# Differential Patterns of Gut and Oral Microbiomes in Hispanic Individuals with Cognitive Impairment

**DOI:** 10.1101/2024.07.27.605455

**Authors:** Yannick N. Wadop, Erin L. Vasquez, Julia J. Mathews, Jazmyn A. S. Muhammad, Rosa Pirela Mavarez, Claudia Satizabal, Mitzi M Gonzales, Jeremy Tanner, Gladys Maestre, Alfred N. Fonteh, Sudha Seshadri, Tiffany F. Kautz, Bernard Fongang

## Abstract

Alzheimer’s disease and related dementias (ADRD) have been associated with alterations in both oral and gut microbiomes. While extensive research has focused on the role of gut dysbiosis in ADRD, the contribution of the oral microbiome remains relatively understudied. Furthermore, the potential synergistic interactions between oral and gut microbiomes in ADRD pathology are largely unexplored. This study aims to evaluate distinct patterns and potential synergistic effects of oral and gut microbiomes in a cohort of predominantly Hispanic individuals with cognitive impairment (CI) and without cognitive impairment (NC). We conducted 16S rRNA gene sequencing on stool and saliva samples from 32 participants (17 CI, 15 NC; 62.5% female, mean age = 70.4 ± 6.2 years) recruited in San Antonio, Texas, USA. Correlation analysis through MaAslin2 assessed the relationship between participants’ clinical measurements (e.g., fasting glucose and blood cholesterol) and their gut and saliva microbial contents. Differential abundance analysis evaluated taxa with significant differences between CI and NC groups, and alpha and beta diversity metrics assessed within-sample and group compositional differences. Our analyses revealed no significant differences between NC and CI groups in fasting glucose or blood cholesterol levels. However, a clear association was observed between gut microbiome composition and levels of fasting glucose and blood cholesterol. While alpha and beta diversity metrics showed no significant differences between CI and NC groups, differential abundance analysis revealed an increased presence of oral genera such as *Dialister*, *Fretibacterium*, and *Mycoplasma* in CI participants. Conversely, CI individuals exhibited a decreased abundance of gut genera, including *Shuttleworthia*, *Holdemania*, and *Subdoligranulum*, which are known for their anti-inflammatory properties. No evidence was found for synergistic contributions between oral and gut microbiomes in the context of ADRD. Our findings suggest that similar to the gut microbiome, the oral microbiome undergoes significant modifications as individuals transition from NC to CI. Notably, the identified oral microbes have been previously associated with periodontal diseases and gingivitis. These results underscore the necessity for further investigations with larger sample sizes to validate our findings and elucidate the complex interplay between oral and gut microbiomes in ADRD pathogenesis.

## Introduction

The intricate interplay between the human microbiome and health has garnered significant attention, with emerging research elucidating its profound influence on physiological functions and disease states.^1–3^ The gut microbiome stands out as a pivotal player among the myriad microbial ecosystems inhabiting the human body.^4–6^ The concept of the gut-brain axis has emerged as a fundamental framework elucidating the bidirectional communication between the gut microbiota and the central nervous system, implicating its involvement in various neurological disorders, including Alzheimer’s disease and related dementias (ADRD).^7–14^

While gut dysbiosis has been extensively studied in the context of ADRD, the role of the oral microbiome, particularly the saliva microbiome, remains elusive despite its potential significance. The oral cavity harbors a diverse microbiota, second only to the gut, comprising over 770 bacterial species.^15,16^ Emerging evidence suggests that the effects of oral dysbiosis could extend beyond oral health, impacting brain health through mechanisms such as the production of inflammatory factors like lipopolysaccharides (LPS), which may facilitate neuroinflammation and neurodegeneration.^17–28^

Environmental factors have been implicated in shaping the oral microbiome, potentially affecting inflammatory responses.^29–34^ Additionally, oral-to-gut and gut-to-oral microbial transmissions can modulate the microbial ecosystems in both habitats, influencing disease pathogenesis. Recent research has begun to shed light on the oral microbiome’s involvement in neurological health, proposing the concept of an oral-brain axis.^19–21,35–37^ For instance, oral dysbiosis in individuals with poor cardiovascular health and/or cognitive impairment is associated with an increase in bacterial genera such as *Tannerella*, *Porphyromonas*, *Dialister*, *Treponema*, *Fretibacterium*, and *Fusobacterium*.^21,23,38,39^ Moreover, oral dysbiosis has been shown to induce inflammatory responses through elevated levels of cytokines, C-reactive protein, white blood cells, and intercellular adhesion molecules.^38,40–43^

Synergistic interactions between the gut and saliva microbiomes have also been reported, underscoring their collective impact on neurological function and disease processes. Indeed, evidence indicates that saliva microbes entering the gastrointestinal tract may disrupt the balance of the gut microbiota and disturb the homeostasis of the immune response in the intestine.^21,35,44,45^

This study aims to investigate the differential patterns of the gut and saliva microbiomes in a cohort of predominantly Hispanic/Latino participants comprising cognitively normal individuals and those with cognitive dysfunction in San Antonio, Texas, USA. By examining the intricate relationships between the gut-brain axis, the oral-brain axis, and brain health, this research seeks to contribute to our understanding of the roles of the human oral and gut microbiomes in neurodegenerative diseases.

## 2. Materials and Methods

### Study population and sample collection

This study was conducted at the University of Texas Health Science Center at San Antonio (UTHSCSA). Questionnaires were used to collect demographic details such as age and sex, as well as pertinent medical history, current medication usage, dietary patterns, and alcohol consumption habits. Participants were excluded if they had used antibiotics in the last 30 days or were younger than 55. Only participants who provided both stool and saliva samples and had recently undergone a comprehensive physical/neurological exam and cognitive assessment using the Uniform Data Set version 3 (USD-3^45–47^) were included in this analysis. Participants with MCI or dementia were clinically diagnosed using the National Institute on Aging-Alzheimer’s Association criteria^48^ and had a Global Clinical Dementia Rating (CDR)^49^ Scale score of 0.5 or 1. Stool and saliva samples were collected by participants at home using OMNIgene ORAL and OMNIgene GUT collection devices (DNAgenotek, Ottawa, ON), shipped to the South Texas Alzheimer’s Disease Research Center (STAC) Biomarker laboratory in San Antonio, TX, and processed/stored according to the manufacturer’s guidelines.

Ethical approval for this study was obtained from the Institutional Review Board of the UTHSCSA, ensuring adherence to ethical principles and guidelines governing human research participants. All procedures involving human participants were conducted in accordance with the ethical standards outlined in the Declaration of Helsinki and its subsequent amendments. Prior to enrollment, participants provided signed, written informed consent indicating their willingness to participate.

### Clinical labs

Standard biological, clinical lab measurements, such as HbA1c, fasting glucose, lipid panels, homocysteine, hsCRP, and creatinine, were performed by a local Labcorp (Burlington, NC). Measurements were not available for all participants. HbA1c (n=32), fasting glucose (n=32), lipid panels (n=32), homocysteine (n=27), hsCRP (n=27), and creatinine (n=31) and were collected on average within 25.6 days (SD=39.7, Min=0, Max=196) of the stool and saliva samples used for microbiome sequencing.

### 16S rRNA gene sequencing analysis and microbial profiling

Stool and saliva samples were collected by participants at home using OMNIgene ORAL and OMNIgene GUT collection devices (DNAgenotek, Ottawa, ON), shipped to the STAC Biomarker laboratory, and stored according to the manufacturer’s guidelines. Bacterial DNA was extracted using the Maxwell RSC Fecal Microbiome kit (Madison, WI). The UTHSCSA Genomic Sequencing Facility performed 16S rRNA (V4 region) gene sequencing (primer 515F:5’-AATGATACGGCGACCACCGAGATCTACACTATGGTAATTGTGTGCCAGCMGCCGCGGTAA-3’) and unique reverse barcode primers from the Golay primer set.^50–52^ After amplification, sample replicates were pooled, cleaned to remove residual contaminants, and sequenced at the UTHSCSA Genomic Sequencing Facility using the Illumina HiSeq 3000. The reads generated from the 16S rRNA gene sequencing were processed and analyzed using the Quantitative Insights Into Microbial Ecology 2 (*QIIME2*) software.^53^ Sequencing reads were clustered into Amplicon Sequencing Variants (ASVs) to delineate individual bacterial taxa present within the samples. To ensure robust and reliable analysis, ASVs with less than four reads in less than 10% of samples were removed to minimize the impact of low representation and sequencing depth variations across samples. Taxonomic classification was performed on the clustered ASVs to assign taxonomy to the sequences, enabling the identification of bacterial taxa at various levels of phylogenetic resolution. Phylogenetic trees were constructed using the *MAFFT* algorithm implemented within the *QIIME2* pipeline.

### Predictive Functional Content of Microbial Communities

To explore the potential functional attributes of the microbial communities within the cohort, we used the Phylogenetic Investigation of Communities by Reconstruction of Unobserved State (*PICRUSt*).^54^ Functional prediction was performed using Kyoto Encyclopedia of Genes and Genomes (KEGG) orthologs (KO), Enzyme Commission (EC) classifications, and MetaCyc metabolic pathways (MePath) enrichments. Differences between the predicted functional contents of cognitively normal (**NC**) and impaired (**CI**) participants were conducted using the *MaAslin2* package.^55^

### Differential Abundance Analysis of Microbial Communities

We analyzed the association between clinical laboratory measures and gut/oral microbial genera using univariable linear regression via *MaAslin2*. The following model was employed to identify genera associated with clinical laboratory measures: *Genera ∼ clinical labs measurements*, where “*Genera*” denotes the relative abundance counts at the genus level. For *MaAslin2* processing, thresholds were set with a minimum relative abundance of >10^-3^ and a minimum detection threshold of 10% of samples for each feature. Associations were considered significant if the Benjamini-Hochberg (BH) adjusted p-value was less than 0.05.

Sample α-diversity was assessed using Chao1, ACE, Observed, Shannon, and Simpson indices. At the same time, β-diversity was visualized through principal coordinates analysis (PCoA) based on Bray-Curtis distances to evaluate group separation in compositional data. Differences in community composition between cognitively impaired (**CI**) and cognitively normal (**NC**) groups were tested using permutational analysis of variance (PERMANOVA) implemented in the vegan package.^56^

Differential abundance (DA) analysis, performed with the R package *MicrobiotaProcess*, identified gut and saliva microbial features that differed between NC and CI groups. DA results were considered significant if they had adjusted p-values <0.05 and a logarithmic linear discriminant analysis (LDA) score >2. We used to perform differential Kyoto Encyclopedia of Genes and Genomes (KEGG) Orthology (KOs) representations between NC and CI participants. Statistically significant KOs (adjusted p-value <0.05) were subsequently subjected to pathway enrichment analysis using the *ClusterProfiler* package in R and visualized using *EnrichPlot*.^57^

## 3. Results

This study included 32 predominantly Hispanic/Latino (%Hispanic = 90.6) participants (%F = 62.5, mean age = 70.4 ± 6.2), of which 17 were classified as CI and 15 were classified as NC. There was no statistically significant differences in age, gender distribution, body mass index or laboratory assays between the CI and NC groups. Detailed demographic and clinical data for all participants are summarized in **Table 1**, providing a comprehensive overview of the study cohort’s characteristics.

**Table 1.**
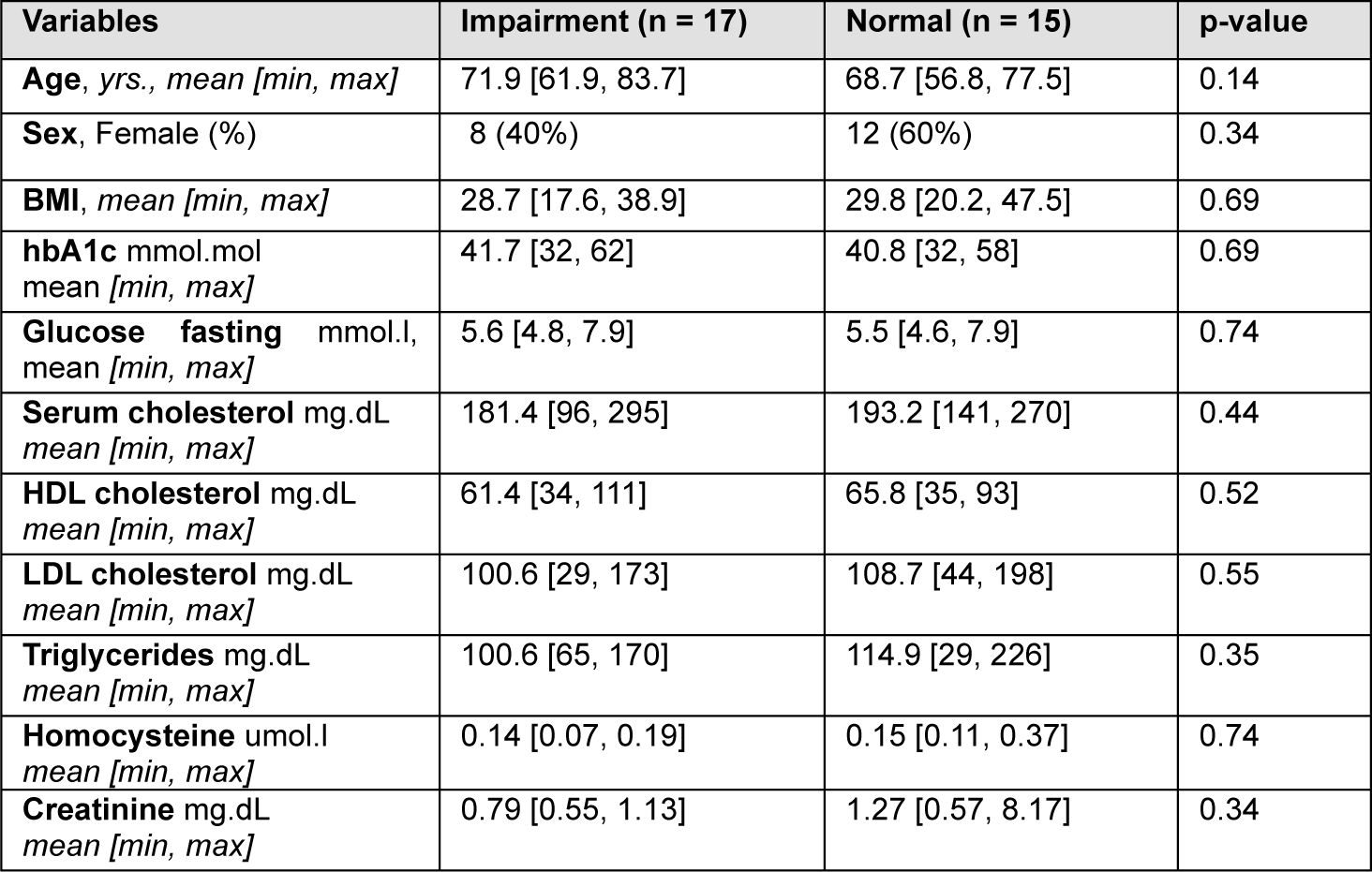
Demographic, BMI, fasting blood glucose and lipids, and markers inflammation, diabetes, and metabolism of cognitively normal and impaired Hispanic participants (%Hispanics/Latinos = 90.6).

| Variables | Impairment (n = 17) | Normal (n = 15) | p-value |
| --- | --- | --- | --- |
| Age, yrs., mean [min, max] | 71.9 [61.9, 83.7] | 68.7 [56.8, 77.5] | 0.14 |
| Sex, Female (%) | 8 (40%) | 12 (60%) | 0.34 |
| BMI, mean [min, max] | 28.7 [17.6, 38.9] | 29.8 [20.2, 47.5] | 0.69 |
| hbA1c mmol.mol<br>mean [min, max] | 41.7 [32, 62] | 40.8 [32, 58] | 0.69 |
| Glucose fasting mmol.l,<br>mean [min, max] | 5.6 [4.8, 7.9] | 5.5 [4.6, 7.9] | 0.74 |
| Serum cholesterol mg.dL<br>mean [min, max] | 181.4 [96, 295] | 193.2 [141, 270] | 0.44 |
| HDL cholesterol mg.dL<br>mean [min, max] | 61.4 [34, 111] | 65.8 [35, 93] | 0.52 |
| LDL cholesterol mg.dL<br>mean [min, max] | 100.6 [29, 173] | 108.7 [44, 198] | 0.55 |
| Triglycerides mg.dL<br>mean [min, max] | 100.6 [65, 170] | 114.9 [29, 226] | 0.35 |
| Homocysteine umol.l<br>mean [min, max] | 0.14 [0.07, 0.19] | 0.15 [0.11, 0.37] | 0.74 |
| Creatinine mg.dL<br>mean [min, max] | 0.79 [0.55, 1.13] | 1.27 [0.57, 8.17] | 0.34 |

### Participants’ oral and gut microbial contents differ between the cognitively impaired and normal groups

We assessed the association between participants’ clinical laboratory measurements and their gut and saliva microbial compositions, operating under the hypothesis that there exists a correlation between physiological parameters and the gut microbiome (**Figure 1A-B**). Through regression analyses, we identified multiple oral and gut bacterial genera significantly associated with two or more clinical laboratory measures (BH adjusted p-value < 0.05) (**Figure 1A-B**). These findings suggest distinct patterns of association between the participants’ physiological states and their gut and oral microbial compositions.

**Figure 1.**
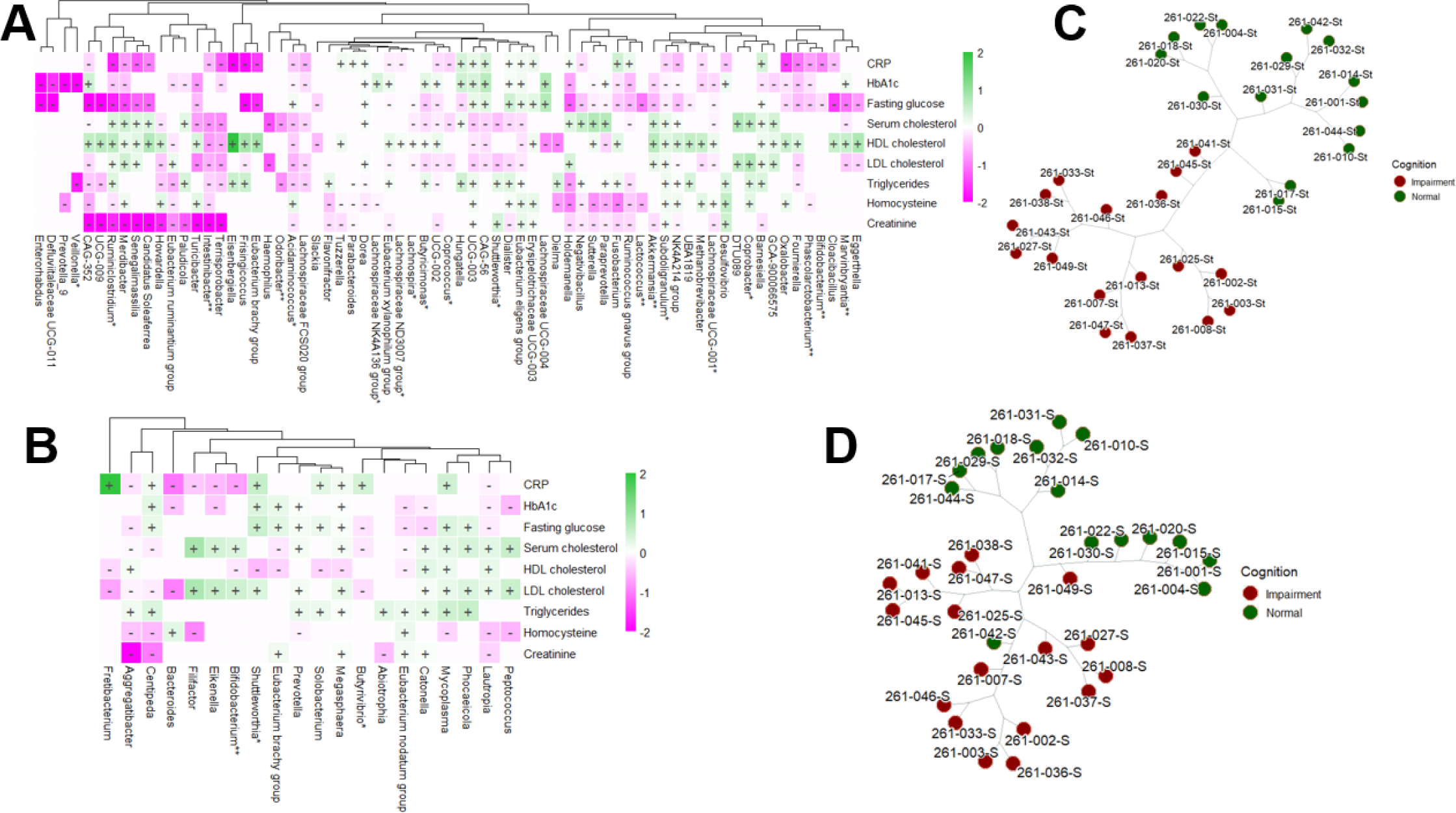
Univariable association analysis using MaAslin2 displaying the positive (+) and negative (-) correlations (BH adjusted p-value < 0.05) between participants’ CRP, hbA1c, fasting blood glucose, blood lipids, homocysteine and creatinine and gut (A) and oral (B) microbiomes. The genera known to be able to produce short-chain fatty acid (SCFA) are marked with an asterisk (*) and those that are probiotics for SCFA are marked with two asterisks (**). Unsupervised phylogenic clustering of gut (C) and oral microbiomes (D) distinguished normal participants from cognitively impaired.

### Fasting glucose and the microbiome

We observed a general negative correlation trend between gut bacterial genera and fasting glucose levels. Specifically, genera such as *Marvinbryantia*, *Bifidobacterium*, *Phascolarctobacterium*, *Oxalobacter*, *Subdoligranulum*, *Butyricimonas*, and *Lachnospira* exhibited these negative associations. Among the 45 correlations examined between bacterial genera and fasting glucose, only 9 bacterial genera showed positive correlations, notably including *Barnesiella*, *Dialister*, *Dorea*, and members of the *Eubacterium eligens* group (refer to **Figure 1A** and **Table SF2** for details). The comprehensive list of gut bacterial genera associated with fasting blood glucose levels is provided in **Table 2**. Additionally, our analysis revealed 13 correlations between saliva microbiome composition and fasting glucose levels. Of these, 5 correlations were negative, involving oral bacterial genera such as *Aggregatibacter*, *Butyrivibrio*, and *Lautropia*. Conversely, 8 correlations were positive, including genera such as *Mycoplasma*, *Prevotella*, and *Megasphaera* (see **Figure 1B**).

**Table 2.**
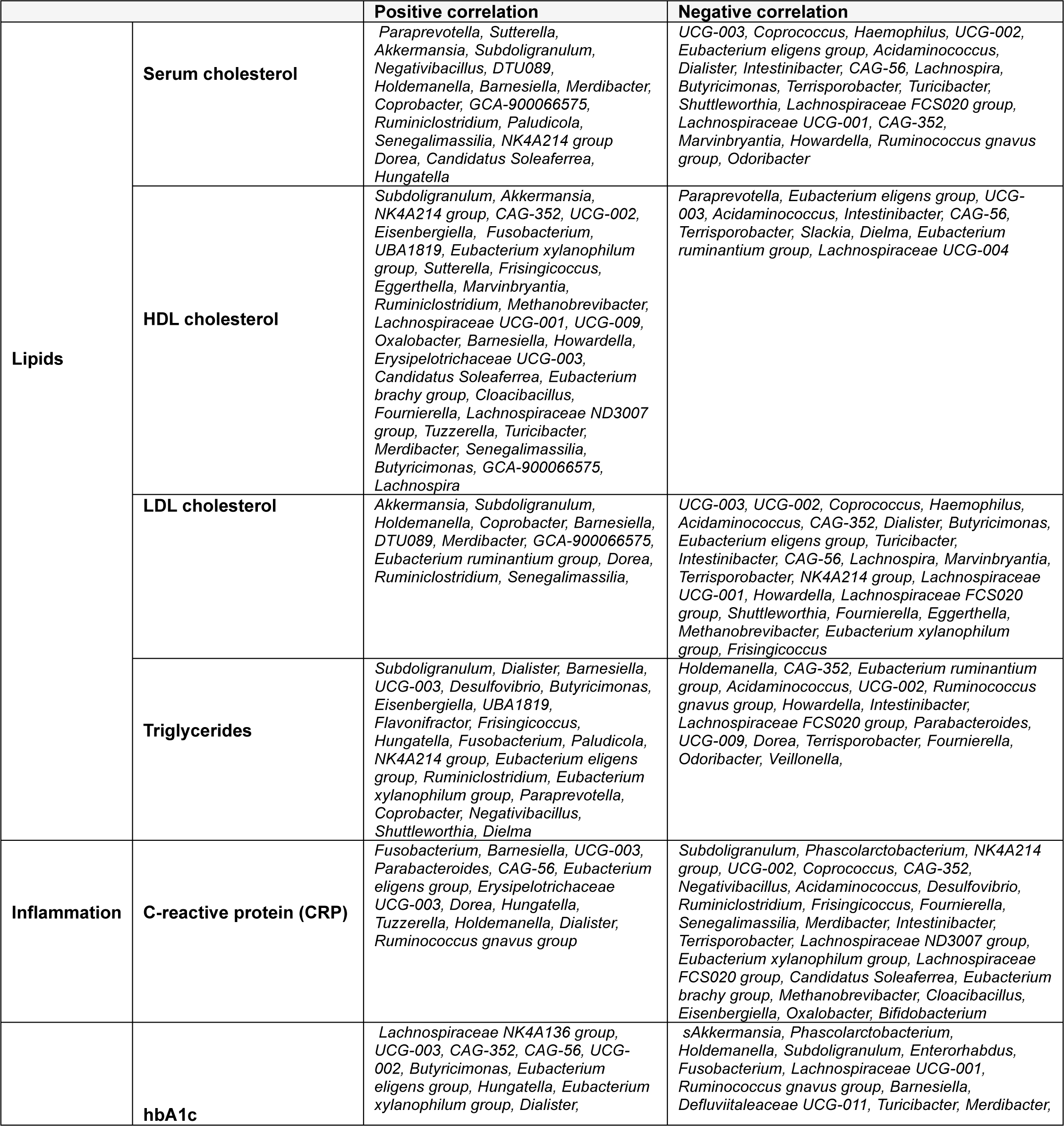
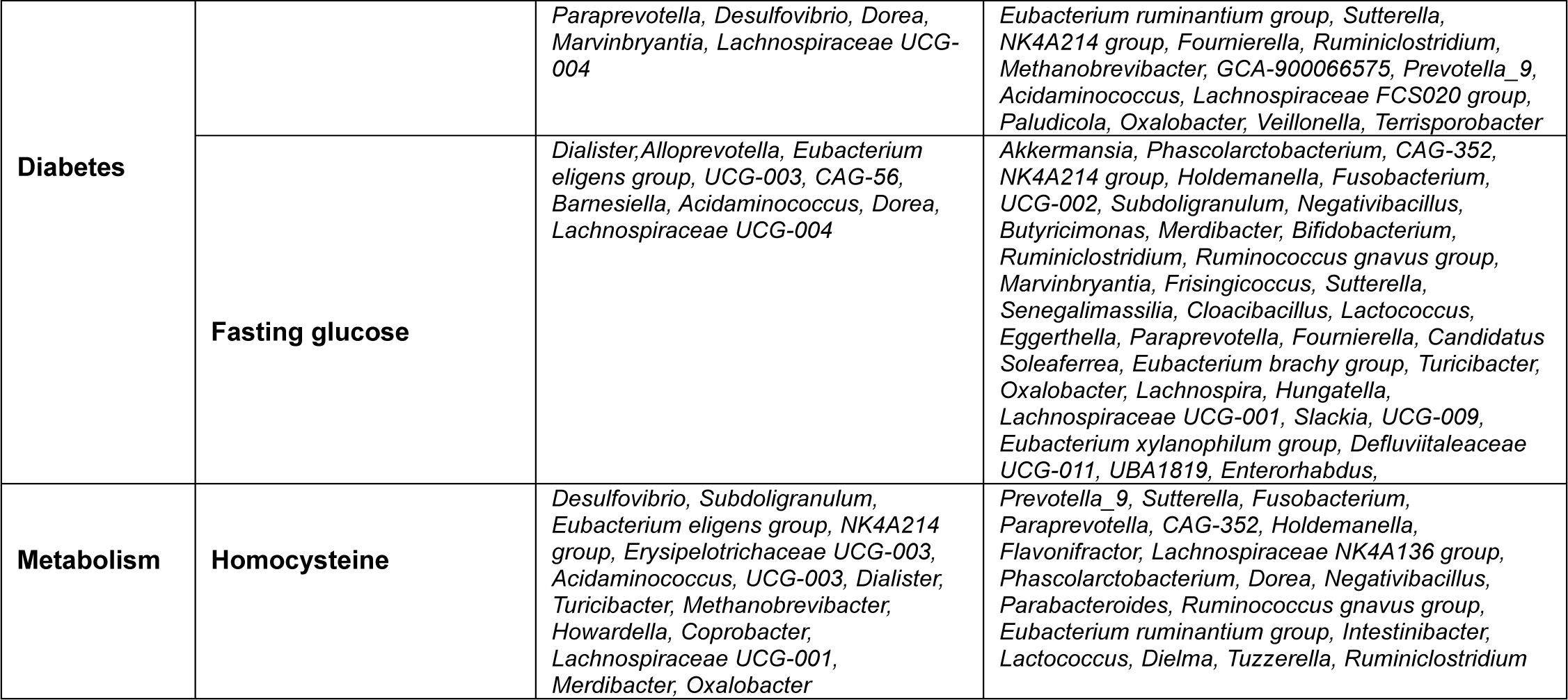
Univariable association of gut bacterial genera and clinical labs measures using MaAslin2. Gut genera showing significant (BH adjusted p-value < 0.05) positive or negative correlation with fasting blood glucose, blood lipids, a marker of systemic inflammation (CRP), diabetes (hbA1c), and overall metabolism (homocysteine).

### Implications of the microbiome on diabetes (hemoglobin A1c)

Our analysis revealed that the majority of gut bacterial genera exhibited negative correlations with hemoglobin A1c (HbA1c) levels. Specifically, out of 41 statistically significant correlations, 26 gut genera demonstrated significant associations with HbA1c. Notable genera among these include *Barnesiella*, *Phascolarctobacterium*, *Oxalobacter*, *Subdoligranulum*, *Akkermansia*, *Methanobrevibacter*, *Veillonella*, and *Prevotella_9* (**Figure 1A** and **Table 2**). In contrast, only 11 oral bacterial genera were found to be significantly correlated with HbA1c levels. The comprehensive list of these correlations is detailed in Supplementary **Table SF2**.

### Fasting blood cholesterol and the microbiome

We observed statistically significant correlations between gut bacterial genera and high-density lipoprotein (HDL) cholesterol, as illustrated in **Figure 1A** and detailed in **Table 2**. Notably, the genera *Oxalobacter, Subdoligranulum, Akkermansia, Barnesiella,* and *Lachnospira* demonstrated positive associations with HDL cholesterol levels. In contrast, oral bacterial genera showed predominantly negative correlations (63.6% of negative correlations) with HDL cholesterol, with fewer instances of positive associations. Furthermore, serum cholesterol and low-density lipoprotein (LDL) cholesterol also exhibited correlations with gut microbiota. Most correlations were negative (60.5% of positive correlations), although some positive associations were observed. The gut bacterial genera showing significant negative correlations with both serum cholesterol and LDL cholesterol included *Marvinbryantia, Shuttleworthia, Coprococcus, Butyricimonas, Lachnospira,* and *Acidaminococcus*. Conversely, oral bacterial genera exhibited mostly positive correlations (67.7% of positive correlations) with serum cholesterol and LDL cholesterol levels, with notable genera including *Bifidobacterium, Peptococcus, Mycoplasma, Prevotella,* and *Megasphaera.* A comprehensive summary of these correlations is presented in Supplementary **Table SF2** and **Table 2**.

### Implications of the microbiome on inflammation (C-reactive protein) and oxidation (homocysteine)

No statistically significant differences in C-reactive protein (CRP) or homocysteine levels were observed between cognitively normal (NC) and cognitively impaired (CI) older adults. However, several gut bacterial genera demonstrated notable correlations with CRP and homocysteine. Specifically, CRP levels were predominantly negatively (66.7% of negative correlations) correlated with various gut genera, including *Coprococcus, Acidaminococcus, Bifidobacterium, Oxalobacter, Marvinbryantia,* and *Subdoligranulum*. In contrast, the correlations between homocysteine levels and gut genera were more evenly distributed, with both positive and negative associations observed. A comprehensive summary of these associations is presented in **Table 2**. Additionally, only a limited number of associations were identified between oral bacterial genera and CRP and homocysteine levels (**Figure 1B** and **Table SF1**).

Subsequently, we performed phylogenetic clustering of the samples to evaluate whether microbial abundance could independently differentiate CI participants from NC (refer to **Figure 1C-D**). Our analyses demonstrated distinct clustering patterns based on the compositions of both the gut (**Figure 1C**) and saliva (**Figure 1D**) microbiomes among the participants.

Altogether, these findings underscore the potential importance of gut microbiota in metabolic health, particularly in the context of glucose metabolism, among individuals with cognitive impairment.

### 3.1. No difference in alpha and beta diversity metrics

To evaluate the diversity of gut and oral microbiomes in NC and CI participants, we utilized Chao1, Observed Species, and Abundance-based Coverage Estimator (ACE) indices to assess species richness, while the Shannon diversity index was employed to estimate both species evenness and richness. Our analysis revealed no significant differences in gut alpha diversity between the NC and CI groups, as depicted in **Figures 2A** and **2B**. Additionally, gut beta diversity, analyzed using Principal Coordinates Analysis (PCoA) with Bray-Curtis distances, showed no significant differences in species composition between the two groups after performing PERMANOVA. Similarly, alpha and beta diversity analyses of saliva samples indicated no significant differences between the NC and CI groups, as shown in **Figures 2C** and **2D**.

**Figure 2.**
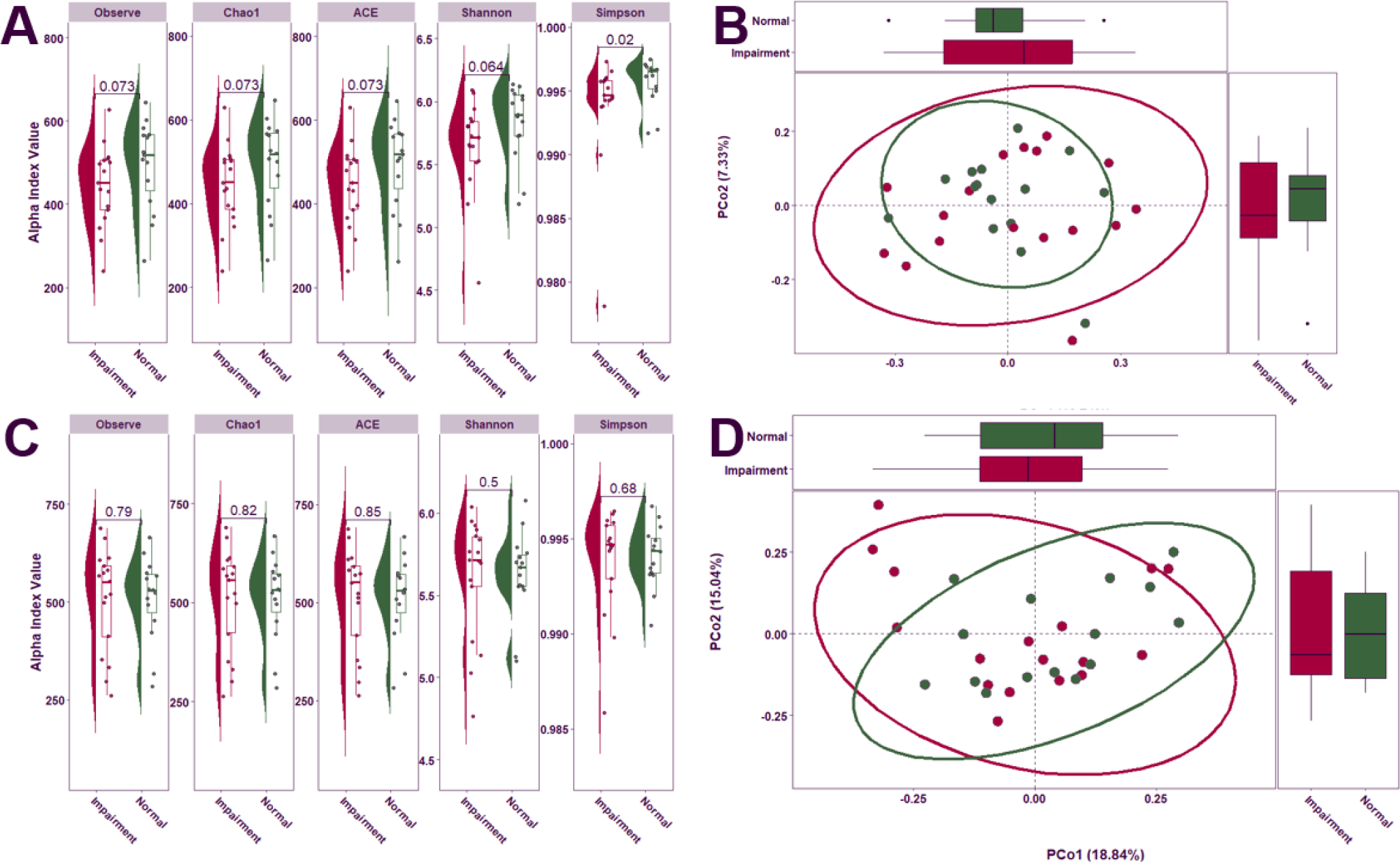
Comparison of the microbial community diversity of samples collected in individuals with CI and NC. Alpha diversity of stool (A) and saliva (C) samples based on 16S rRNA sequencing. The tests of difference in microbial diversity (Chao1, ACE, Observe, and Shannon indexes) at the OTU level between CI and NC were based on Wilcoxon test. Beta diversity on stool (B) and saliva (D) samples represented by PCoA with Bray-Curtis distances. No differences were observed between groups based on the PERMANOVA test (P = 0.83).

### 3.2. Oral genera are differentially abundant between NC and CI participants

The differential abundance analysis of oral microbiome composition revealed notable shifts in several genera among CI individuals. Specifically, there was a marked decrease in the relative abundance of *Neisseria* and an increase in *Streptococcus* and *Fusobacterium* in CI participants compared to those who were cognitively normal (**Figure 3A**). For biomarker discovery, we employed Linear Discriminant Analysis (LDA) to assess the effect sizes and relative abundances driving these differences. LDA effect sizes indicate the magnitude of consistent differences in relative abundance between features (bacterial taxa) in CI and NC individuals. Using this approach, we identified 13 taxonomic biomarkers that were significantly overrepresented in CI cases (adjusted p-value < 0.05, with LDA effect sizes > 2; **Figure 3B and Table ST2**).

**Figure 3.**
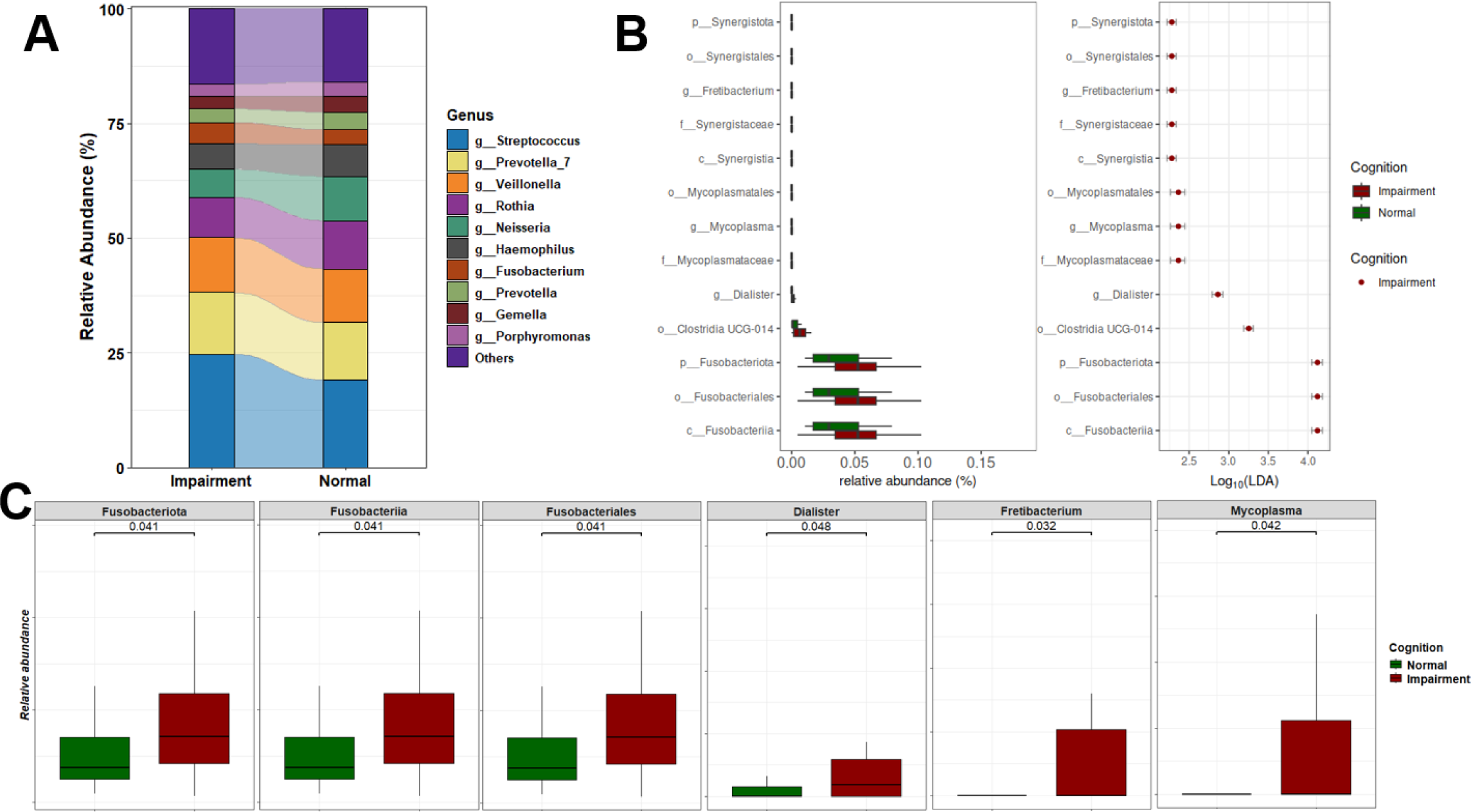
(A) Stacked plot illustrating the microbiome profiles of predominant taxa in the oral microbiomes of CI and NC participants at the genus level. (B) LDA results on saliva microbiome detected 13 bacterial taxa showing statistically significant and biologically consistent differences in CI at the phylum (p-), class (c-), order (o-), family (f-), and genus (g-) levels. (C) Relative abundance of the most differential taxa and identified oral genera between CI and NC participants.

Among the top differentially abundant taxa were *Fusobacteriia*, *Fusobacteriota*, and *Fusobacteriales*. We also observed a significant increase in the abundance of several genera in CI participants, including *Fretibacterium*, a gram-negative bacterium previously linked to periodontitis; *Mycoplasma*, a gram-negative species prevalent in individuals with gingivitis and periodontitis; and *Dialister*.^58,59^ These bacteria are commonly associated with oral and systemic diseases, particularly periodontitis, gingivitis, and advanced periodontal conditions.^60,61^ Furthermore, the phyla *Synergistota* and *Fusobacteria* were significantly increased in CI participants. **Figure 3C** highlights the top statistically significant taxa, underscoring their varying abundance between NC and CI groups.

### 3.3. Differential Abundance analysis of the Gut Microbiome in CI and NC participants

Significant differences emerged in the gut microbiome composition between individuals with normal cognition and those with cognitive impairment. As depicted in **Figure 4A**, the relative abundances of gut genera varied markedly between the two groups. Specifically, individuals with CI exhibited an increased abundance of *Bacteroides* and *Escherichia-Shigella*, and a decreased abundance of *Ruminococcus* and *Subdoligranulum*. Further analysis utilizing Linear Discriminant Analysis (LDA) identified several taxonomic biomarkers with significantly lower abundances in the CI group (**Figure 4B** and **Table ST1**). Notably, genera known for their anti-inflammatory functions, such as *Subdoligranulum*, *Holdemania*, *Shuttleworthia*, UGC-005, and the *Eubacterium brachy* group, were found to be reduced in CI individuals. Additionally, *Clostridia*, *Firmicutes*, and *Oscillospirales* were the most differentially abundant gut bacteria, with effect sizes of 4.61, 4.54, and 4.49, respectively. Moreover, a significant reduction in the overall abundance of the *Firmicutes* phylum was observed in CI participants (**Figure 4C**). This depletion further underscores the complex dysbiosis present in individuals with cognitive impairment, highlighting the potential functional implications of these microbial alterations on cognitive function.

**Figure 4.**
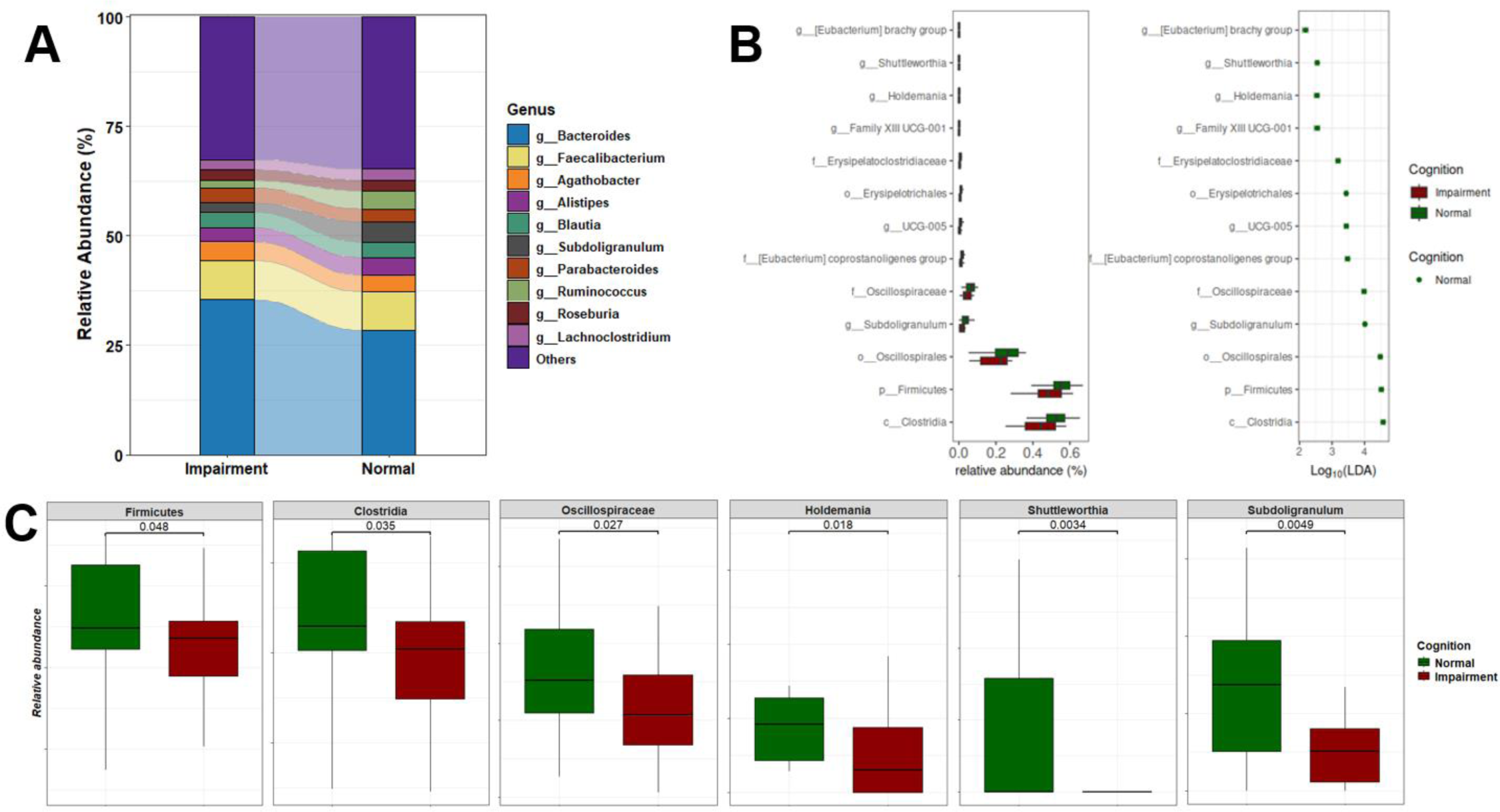
(A) Stacked plot illustrating the microbiome profiles of predominant taxa in the gut microbiome of CI and NC participants at the genus level. (B) LDA results on gut microbiome detected 13 bacterial taxa showing statistically significant and biologically consistent differences in CI at the phylum (p-), class (c-), order (o-), family (f-), and genus (g-) levels. (C) Relative abundance of the most differential taxa and identified gut genera between CI and NC participants.

### 3.4. Functional potential of the gut and oral microbial communities associated with cognitive health

We employed Phylogenetic Investigation of Communities by Reconstruction of Unobserved States (*PICRUSt*) analysis to assess the functional potential of the gut microbiota differentially represented in CI and NC individuals.

This analysis revealed 487 predicted metabolic pathways (MePath), 2,913 predicted enzyme commission numbers (ECs), and 10,543 predicted Kyoto Encyclopedia of Genes and Genomes orthologs (KOs). Subsequent differential analysis identified 33 MePath that exhibited significant differences (BH adjusted p-value < 0.05) between CI participants and NC individuals. Among these differential pathways, several are known to play crucial roles in host metabolism, including glucose degradation, glycol metabolism and degradation, glycolysis, pyruvate dehydrogenase, the tricarboxylic acid (TCA) cycle, glyoxylate bypass, glyoxylate cycle, phospholipases, TCA cycle IV (2-oxoglutarate decarboxylase), photorespiration, fatty acid biosynthesis (E. coli), and L-glutamate degradation VIII (to propanoate). Notably, with the exception of glutamate degradation VIII, these pathways were enriched in CI participants compared to NC individuals (see **Figure SF1A**).

In our analysis, we identified 14 oral metabolic pathways (MePath) that were significantly altered between CI and NC participants (see **Figure SF1B**). Notably, pathways such as ketogluconate metabolism, enterobacterial common antigen biosynthesis, 4-hydroxyphenylacetate degradation, and aerobactin biosynthesis were found to be significantly reduced in CI participants. Conversely, the pathways involved in L-glutamate degradation VIII and L-tryptophan biosynthesis were elevated in CI participants. Further examination of enzyme commission (EC) terms revealed 160 distinct gut EC terms and 120 oral EC terms with significant differences between CI and NC participants (see **Table ST2**). Additionally, our analysis of gene orthologs (KOs) indicated that 1,025 KOs were differentially expressed in the gut microbiome between NC and CI participants, while 616 KOs showed differential expression in saliva samples (see **Table ST3**). To elucidate the biological relevance of these findings, we employed a pathway enrichment analysis using the differentially expressed KOs to map the activated KEGG pathways associated with the transition from NC to CI. This approach provides a more nuanced understanding of the functional roles of the gut and saliva microbiomes in cognitive health.

#### 3.4.1 KEGG pathways associated with gut microbiome profiles in cognitive health

We aimed to explore the biological pathways associated with the differentially expressed KEGG Orthologs (KOs) identified between CI and NC individuals. We identified 28 KEGG pathways that were significantly enriched (Benjamini-Hochberg adjusted p-value < 0.05) in the differential KOs between NC and CI participants. These pathways include lipopolysaccharide biosynthesis, bacterial chemotaxis, beta-alanine metabolism, propanoate metabolism, methane metabolism, and bacterial secretion systems. **Figure 5A** illustrates the number of differential gut KOs associated with each enriched pathway, and a comprehensive summary of these pathways is provided in **Table ST4**. Remarkably, many of these KEGG pathways have been previously implicated in cognitive deficits and Alzheimer’s disease and related dementias (ADRD) in both animal models and clinical studies.^62–64^ To further elucidate these associations, we constructed a network diagram (**Figure 5B**) depicting the relationships between differential gut KOs and KEGG pathways. In this network, blue and red nodes represent gut KOs that were upregulated and downregulated in CI individuals, respectively. For example, differential gut KOs such as K02848 (waaP), K03275 (waaO), K03276 (waaR), K07264 (arnT), K11211 (kdkA), K12973 (pagP), K12975 (eptB), K12981 (waaZ), and K12985 (waaW), which are involved in lipopolysaccharide biosynthesis, were found to be upregulated in CI participants. Analysis of these enriched KEGG pathways revealed several notable associations. Methane metabolism was found to correlate with propanoate metabolism, and glyoxylate and dicarboxylate metabolism showed associations with carbon metabolism as well as glycine, serine, and threonine metabolism (see **Figure SF2**). Additionally, beta-alanine metabolism demonstrated a connection with propanoate metabolism. However, no direct associations were observed between lipopolysaccharide biosynthesis and bacterial chemotaxis with other pathways. To provide a spatial representation, we mapped the identified gut KOs involved in beta-alanine metabolism onto a KEGG metabolic map (**Figure SF3**), highlighting these KOs in red.

**Figure 5.**
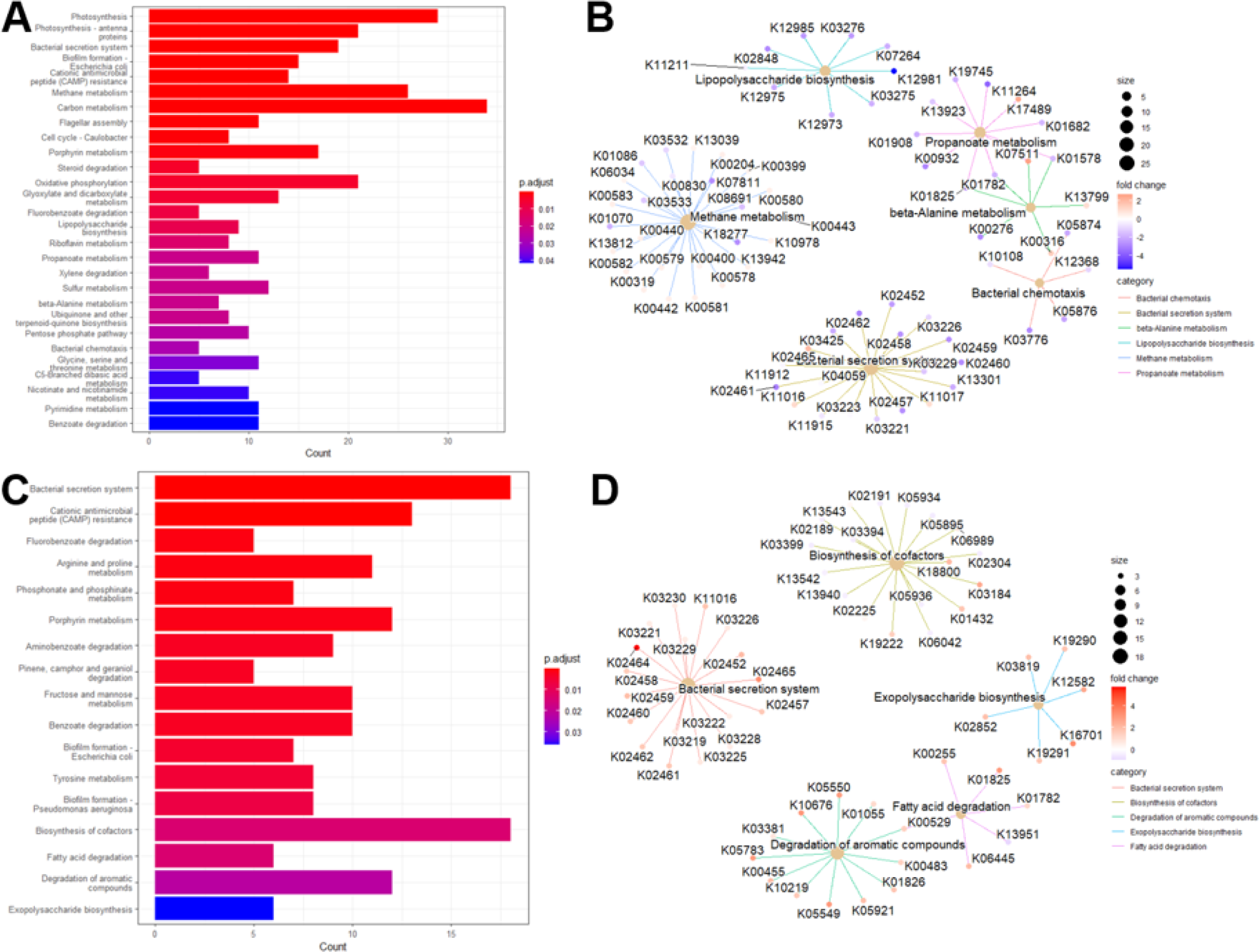
Barplot of enriched KEGG pathways for stool (A) and saliva (C). Network plot of some enriched KEGG pathways displaying the linkages with differential gut KOs for stool (B) and saliva (D).

#### 3.4.2 KEGG pathways associated with oral microbiome profiles in cognitive health

We performed a differential representation analysis of KEGG pathways to elucidate the biological pathways associated with oral KOs and their implications for cognitive health. Our analysis identified 17 KEGG pathways with significant enrichment, including critical pathways such as fatty acid degradation, biosynthesis of cofactors, degradation of aromatic compounds, exopolysaccharide biosynthesis, and the bacterial secretion system. The relationship between these differential oral KOs and the enriched KEGG pathways is depicted in **Figure 5**, with a comprehensive summary provided in **Table ST5**. Notably, **Figure 5D** details the differential oral KOs associated with each enriched KEGG pathway. Among the identified pathways, the bacterial secretion system, biosynthesis of cofactors, cationic antimicrobial peptide (CAMP) resistance, and degradation of aromatic compounds emerged as the top four pathways with the highest counts of differential oral KOs. These findings are consistent with previous research highlighting associations between enriched KEGG pathways and cognitive function, as well as alterations in microbiome composition.^65–69^ Further analysis of the interactions among the identified KEGG pathways (see **Figure SF4**) revealed a strong correlation between Benzoate degradation and the degradation of aromatic compounds, as well as Fluorobenzoate degradation. The degradation of aromatic compounds also exhibited associations with Tyrosine metabolism and Fluorobenzoate degradation. Additionally, connections were observed between Porphyrin metabolism and the biosynthesis of cofactors, and between biofilm formation and the bacterial secretion system. Fatty acid degradation was linked to Pinene, Camphor, and Geraniol degradation. To illustrate these relationships, we mapped the identified oral KOs involved in fatty acid degradation onto a KEGG metabolic map, underscoring their relevance to cognitive health (see **Figure SF5**).

### 3.5. On the synergistic contribution of oral and gut microbiomes to brain health

Previous studies have highlighted a synergistic association between oral and gut microbiomes and brain health. ^39,70^ To further elucidate this relationship, we conducted a detailed examination of the overlap in microbiome taxa and potential functional profiles across these two habitats. Our analysis identified no overlapping differentially abundant taxa between the oral and gut microbiomes in our sampled population. This finding suggests that, within our cohort, there is no apparent synergistic effect in microbial composition between these two sites. To extend our investigation into potential functional synergies, we analyzed KEGG pathways associated with each microbiome. We identified 22 KEGG pathways unique to the gut microbiome, 11 unique to the oral cavity, and 5 pathways that were shared between the two environments (see **Figure 6**). The shared pathways included essential functions such as *Benzoate degradation, Fluorobenzoate degradation, Porphyrin metabolism, Cationic antimicrobial peptide* (CAMP) resistance, biofilm formation in *Escherichia coli*, and bacterial secretion systems. These findings highlight specific functional overlaps that may be relevant to understanding the interactions between the gut and oral microbiomes in relation to cognitive health. Further research is necessary to elucidate the mechanisms underlying these pathways and their potential impacts on cognitive processes, which will advance our understanding of their roles in brain health.

**Figure 6.**
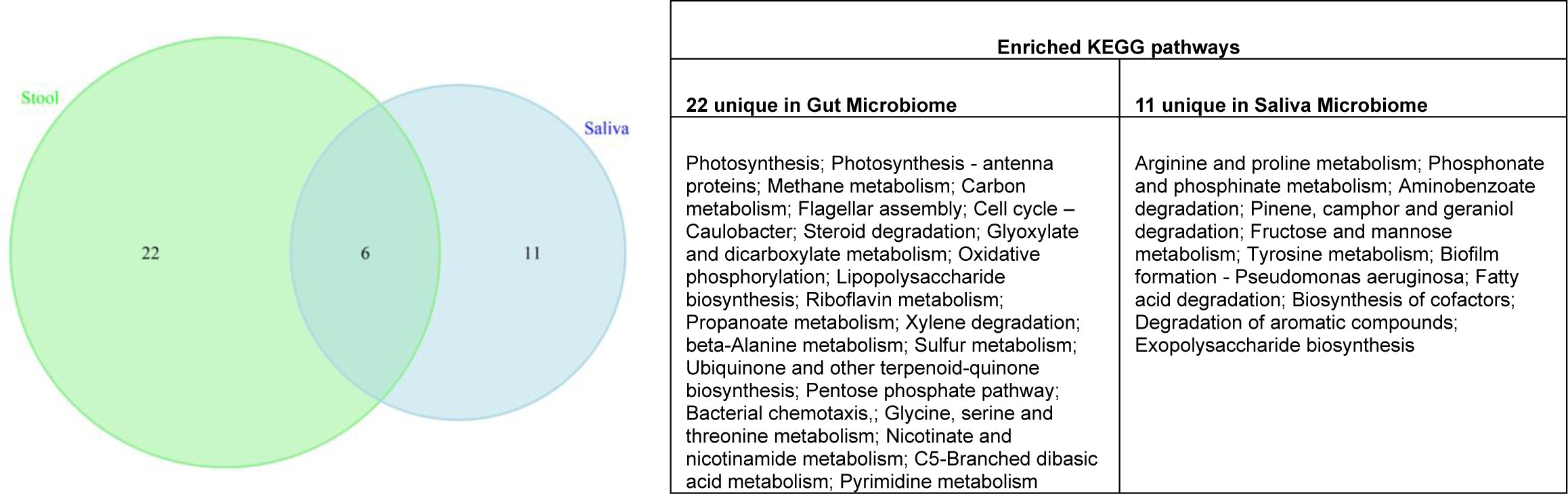
Venn diagram displaying the similarity and overlap KEGG pathways between saliva and gut microbiotas in cognition. These 6 overlapped KEGG pathways included Benzoate degradation, Fluorobenzoate degradation, Porphyrin metabolism, Cationic antimicrobial peptide (CAMP) resistance, biofilm formation in Escherichia coli, and bacterial secretion system.

**Figure 7.**
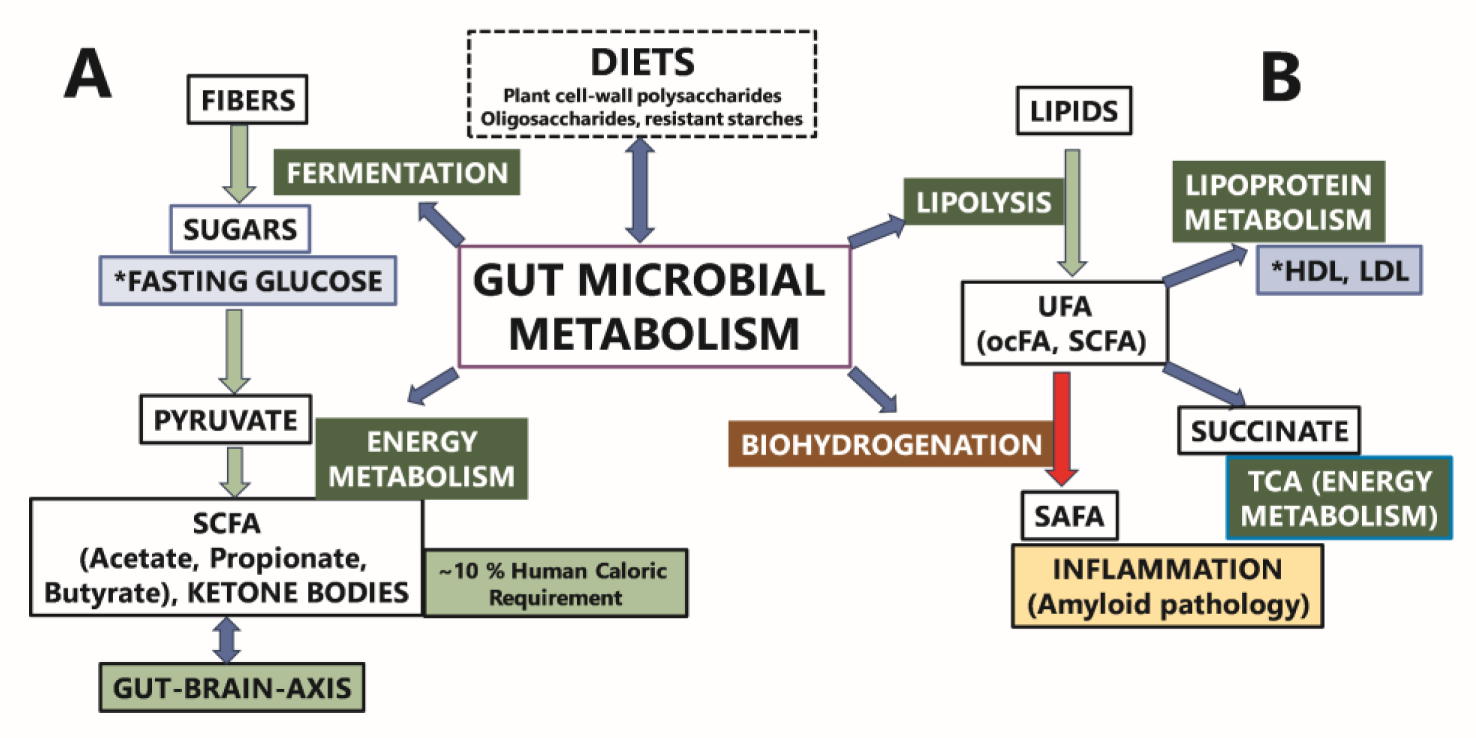
illustrates a proposed interaction between gut microbiome metabolism and the regulation of fasting blood glucose. Complex plant-derived fibers, which are resistant to human digestive enzymes, undergo bacterial fermentation within the gut. This fermentation process converts these complex carbohydrates into hexoses (e.g., glucose) and pentoses, which can then be utilized for energy production through pyruvate and the tricarboxylic acid (TCA) cycle. Short-chain fatty acids, produced by gut microbiome metabolism, play a significant role in colonocyte energy metabolism, contributing approximately 70% of their energy needs. SCFAs diffuse into the bloodstream, where they provide roughly 10% of the total caloric requirement and influence health by modulating various pathways, including immune, endocrine, vagal, and other humoral pathways, such as neurotransmitter regulation. This modulation is a crucial component of the complex gut-brain axis. We hypothesize that dietary fibers and specific gut bacterial genera that regulate glucose levels and support SCFA production are more likely to be associated with cognitive health. Additionally, our findings reveal that gut bacterial genera associated with both fasting glucose and HDL cholesterol levels exhibit opposing directional associations. Overall, distinct bacterial taxa combinations were identified in association with fasting glucose, lipid levels, and plasma metabolites.

## 4. Discussion

The human microbiome plays a critical role in maintaining brain health; however, the specific interactions between alterations in the oral and gut microbiomes within underrepresented populations, particularly Hispanics, are not well elucidated. To address this gap, we examined the shifts in composition and functional profiles of the oral and gut microbiota in individuals with cognitive impairment compared to those with normal cognitive function. Our analysis revealed significant differences in microbial composition at the genus level. Notably, there was an increase in the abundance of taxa with known pathogenic potential in the oral cavity, alongside a reduction in beneficial taxa within the gut microbiota of CI participants. Despite these observed alterations in microbial communities within each habitat, we found no evidence of a synergistic interaction between oral and gut microbiota in relation to brain health.

Although alpha and beta diversity metrics did not reveal statistically significant differences, differential abundance analysis identified notable changes in taxa among participants with cognitive impairment. Specifically, the oral microbiome exhibited a predominance of gram-negative taxa, such as *Dialister*, *Fretibacterium*, and *Mycoplasma*. These taxa have been implicated in inflammatory responses associated with oral diseases, including periodontitis and gingivitis.^71–74^ Recent studies of microbiome data from Hispanic adults have shown that the proliferation of genera like *Dialister*, *Fretibacterium*, and *Mycoplasma* correlates with severe periodontal disease, a known risk factor for Alzheimer’s disease.^75^ Our results align with a previous meta-analysis that examined European and American cohorts, which reported that binge drinking induces a shift in the oral microbiome, characterized by an increased abundance of *Dialister* and *Fretibacterium*, and associates these genera with AD.^76^ Additionally, the enrichment of genera associated with bacteremia in CI participants supports prior research suggesting a potential pathway for bacterial dissemination from the oral cavity to the systemic circulation, possibly affecting brain function.^77^ This hypothesis is corroborated by recent studies demonstrating significant alterations in the oral microbiota profiles of individuals with AD, highlighting the importance of oral microbiota in neurodegenerative conditions.^23,78^ Clinical findings have also linked poor oral hygiene to the proliferation of invasive *Dialister* species, suggesting a potential relationship between oral health and cognitive impairment.^79^ Furthermore, our data reveal positive correlations between *Fretibacterium* and *Mycoplasma* and C-reactive protein levels, a marker of systemic inflammation, indicating a possible overlap between oral microbiota, inflammation, and cognitive health. These findings emphasize the critical role of the oral microbiome in both health and disease, particularly its association with cognitive function.

The analysis of the gut microbiome in individuals with CI revealed a significant decrease in the abundance of beneficial bacteria, particularly within the Firmicutes phylum, which includes key short-chain fatty acid (SCFA) producers.^80^ Notably, *Subdoligranulum*, a Gram-positive bacterium closely related to the *Faecalibacterium* genus, known for its butyrate production, exhibited reduced abundance in CI participants. Butyrate, a SCFA, is increasingly recognized for its potential probiotic effects and its role in improving metabolic health. ^81^ Additionally, *Holdemania*, an acetate-producing bacterium involved in SCFA metabolism, was found to be less prevalent among CI individuals. Higher levels of *Holdemania* have been associated with anti-inflammatory effects and maintenance of gut barrier integrity in both rodent models and humans.^82,83^

Recent investigations have also highlighted the role of *Shuttleworthia*, a Gram-positive, saccharolytic bacterium capable of producing acetate, butyrate, and lactate.^84^ *Shuttleworthia* is essential for sustaining gut health. The observed dysbiosis, characterized by a reduced abundance of these beneficial bacteria, suggests a compromised protective mechanism crucial for maintaining gut barrier integrity. This dysbiosis may contribute to increased gut permeability and subsequent systemic inflammation. Furthermore, the absence of overlapping taxonomic biomarkers between oral and gut microbiomes emphasizes the distinct microbial profiles of these environments. The activation of immune responses and cytokine production in CI could facilitate the translocation of inflammatory agents to the intestine, thereby compromising gut barrier function and potentially exacerbating neuroinflammation and cognitive decline.^85–87^

Despite the lack of significant differences in C-reactive protein levels between cognitively impaired and normal cognition groups, we observed notable correlations between CRP levels and gut microbiota. Specifically, CRP levels were negatively correlated with several gut bacterial genera, including *Bifidobacterium*, *Subdoligranulum*, *Marvinbryantia*, *Acidaminococcus*, and *Coprococcus*. In contrast, associations with oral bacterial genera were less pronounced; however, *Mycoplasma*, *Fretibacterium*, and *Megasphaera* were positively correlated with CRP levels, whereas *Bifidobacterium*, *Aggregatibacter*, and *Bacteroides* exhibited negative correlations.

Hemoglobin A1c and fasting glucose, both critical for diagnosing diabetes, showed distinct correlations with gut microbiota. Negative correlations were observed with genera such as *Subdoligranulum*, *Akkermansia*, *Oxalobacter*, and *Ruminiclostridium*. Conversely, positive correlations were noted with *Dorea*, *Dialister*, and *CAG-56*. Additionally, HbA1c and fasting glucose levels exhibited positive correlations with oral bacterial genera *Shuttleworthia*, *Centipeda*, *Prevotella*, and *Megasphaera*, while negative correlations were observed with *Catonella* and the *Eubacterium nodatum* group. Regarding homocysteine, a marker of oxidative stress, no significant differences were found between NC and CI groups. However, homocysteine levels were positively correlated with gut bacterial genera *Acidaminococcus*, *Coprobacter*, *Subdoligranulum*, and *Oxalobacter*. Homocysteine also showed positive associations with oral genera *Bacteroides* and the *Eubacterium nodatum* group, and a negative correlation with *Mycoplasma*. We did not observe significant differences in lipid profiles or serum metabolites between NC and CI participants. Nonetheless, HDL cholesterol levels demonstrated positive correlations with gut bacterial genera such as *Subdoligranulum*, *Marvinbryantia*, *Butyricimonas*, *Lachnospira*, and *Oxalobacter*. Serum metabolites exhibited more negative correlations with gut genera, including *Shuttleworthia*, *Acidaminococcus*, *Odoribacter*, *Marvinbryantia*, *Coprococcus*, *Butyricimonas*, and *Lachnospira*. Few associations were found between HDL cholesterol levels and oral bacterial genera, with *Fretibacterium* negatively correlated with HDL cholesterol and *Mycoplasma* positively correlated with both serum metabolites and HDL cholesterol. Overall, these findings suggest that gut and oral microbial populations may have distinct functional roles, influencing different physiological and biochemical pathways.

The functional roles of microbial communities were further elucidated through *PICRUSt* analysis, identifying enriched pathways associated with cognitive dysfunction. Notably, lipopolysaccharide-induced neuroinflammation was found to be associated with cognitive impairment through mechanisms that enhance beta-amyloid generation.^62^ Additionally, bacterial chemotaxis may contribute to or exacerbate cognitive impairment, highlighting the adaptive responses of gut microbiota to environmental changes.^63,88^ Furthermore, beta-alanine metabolism was identified as a significant pathway related to cognitive performance, with evidence suggesting that beta-alanine supplementation may improve cognitive function, particularly among older adults with borderline to suboptimal cognitive levels.^62,89,90^ Propanoate metabolism, which has been closely associated with Alzheimer’s disease, was also implicated in disease progression. Elevated propionate levels, potentially resulting from the activity of the Bacteroidetes phylum—the primary producer of propionate in the human gut— could play a role in this process.^63,91^ In contrast, methane metabolism has been recognized for its potential to mitigate spatial memory deficits, likely through the modulation of pro-inflammatory cytokines and microglial activation within the hippocampus.^64,92,93^

Our pathway analysis identified several KEGG pathways relevant to the interaction between the saliva microbiome and cognitive function, as well as alterations in microbiome composition. Notably, perturbations in fatty acid homeostasis and associated pathways have been linked to cognitive impairment and dementia.^94^ The biosynthesis of cofactors pathway has been specifically associated with cognitive deficits in Alzheimer’s disease.^65,66^ Additionally, *cathelicidins*, which are key components of the innate immune response, play a critical role in host defense against microbial infections.^67–69^ The pathway related to exopolysaccharide biosynthesis has been associated with cognitive function enhancement in Alzheimer’s disease, potentially due to its anti-inflammatory effects. Furthermore, our analysis revealed KEGG orthologs involved in lipopolysaccharide biosynthesis in the gut and fatty acid degradation in the oral cavity, suggesting a potential synergistic contribution of both oral and gut microbiome pathways to cognitive impairment. However, it is important to note that the mere overlap of these pathways between the oral and gut microbiomes may not be sufficient to fully support the hypothesized synergistic effects, as previously described.^39,70^

This study offers valuable insights into the relationship between microbial communities and cognitive health; however, several limitations should be noted. Firstly, the relatively small sample size may restrict the generalizability of the findings. Additionally, the dynamic nature of both oral and gut microbiota underscores the need for further longitudinal studies to capture temporal variations and their potential impact on cognitive health. Furthermore, variables such as medication use and dietary intake, known to influence microbial composition, were not accounted for in this study. Addressing these limitations in future research will be essential for a more comprehensive understanding of the complex interactions between microbial communities and cognitive function, as well as for developing targeted therapeutic interventions for neurodegenerative diseases.

## 5. Conclusion

In this study, we analyzed stool and saliva samples to explore the distinct patterns of gut and oral microbiomes in predominantly Hispanic/Latino individuals in San Antonio, Texas, categorized into cognitively normal and cognitively impaired groups. Our findings reveal that cognitive impairment is associated with notable disruptions in both the gut and oral microbiomes. Consistent with existing literature, our results indicate that the oral microbiome in CI participants exhibits altered abundance and functional profiles, with a notable presence of taxa previously linked to periodontal disease and gingivitis. These findings align with growing evidence suggesting a potential bidirectional relationship between oral health and cognitive function. Moreover, our study highlights significant compositional shifts in the gut microbiome of CI participants, particularly a reduction in Firmicutes bacteria known for their anti-inflammatory properties. This observation echoes previous research that underscores the role of gut microbiota in modulating inflammatory pathways and their potential impact on neurodegenerative conditions. Notably, we identified dysregulated pathways in CI individuals, including lipopolysaccharide biosynthesis in the gut and fatty acid degradation in the oral microbiome. These disrupted pathways may contribute to the inflammatory milieu associated with cognitive decline. Our results reinforce the notion that dysbiosis in both the oral and gut microbiomes is intricately linked to cognitive dysfunction, particularly within Hispanic populations. To substantiate these findings, future research should encompass larger and more heterogeneous cohorts to validate these associations and further elucidate the interplay between oral and gut microbiotas in cognitive impairment.

## ACKNOWLEDGMENTS

We thank the many study participants, researchers, and staff for collecting and contributing to the data.

## AUTHOR CONTRIBUTIONS

B.F. conceived and supervised the project, T.K. supervised participants’ recruitment and data collection, E.V., J.M., J.A.S.M. consented participants, collected and processed the samples for sequencing. Y.W. performed all statistical analyses and drafted the results. R.P.M., G.M, and S.S. provided all resources for the project. Finally, Y.W., E.V., J.M., J.A.S.M., R.P.M., G.M., S.S., T.K. and B.F. discussed the results and wrote the manuscript. All authors read and approved the manuscript.

