## Supplementary file for "Differential Patterns of Gut and Oral Microbiomes in Hispanic Individuals with Cognitive Impairment"

### 1. Participants' oral and gut microbial contents differ between the cognitively impaired and normal groups

| | | **Positive correlation** | **Negative correlation** |
|---|---|---|---|
| **Lipids** | **Serum cholesterol** | Prevotella, Eikenella, Lautropia, Filifactor, Megasphaera, Bifidobacterium, Catonella, Peptococcus, Mycoplasma, Phocaeicola | Butyrivibrio, Centipeda, Aggregatibacter, Eubacterium nodatum group, Eubacterium brachy group |
| | **HDL cholesterol** | Catonella, Lautropia, Aggregatibacter, Mycoplasma | Megasphaera, Solobacterium, Eubacterium nodatum group, Eubacterium brachy group, Shuttleworthia, Bifidobacterium, Fretibacterium |
| | **LDL cholesterol** | Eikenella, Prevotella, Megasphaera, Bifidobacterium, Lautropia, Filifactor, Catonella, Shuttleworthia, Peptococcus, Mycoplasma, Phocaeicola | Bacteroides, Butyrivibrio, Fretibacterium, Aggregatibacter, Centipeda |
| | **Triglycerides** | Prevotella, Eubacterium nodatum group, Centipeda, Catonella, Solobacterium, Mycoplasma, Abiotrophia, Aggregatibacter, Phocaeicola, Megasphaera | |
| **Inflammation** | **C-reactive protein (CRP)** | Butyrivibrio, Solobacterium, Fretibacterium, Megasphaera, Shuttleworthia, Mycoplasma, Centipeda, | Eikenella, Bifidobacterium, Aggregatibacter, Bacteroides, Lautropia, Filifactor |

| | | | |
|---|---|---|---|
| **Diabetes** | **hbA1c** | Prevotella, Centipeda, Shuttleworthia, Eubacterium brachy group, Megasphaera | Eubacterium nodatum group, Eikenella, Lautropia, Catonella, Bacteroides, Peptococcus |
| | **Glucose** | Prevotella, Solobacterium, Centipeda, Shuttleworthia, Eubacterium brachy group, Mycoplasma, Megasphaera, Phocaeicola | Eubacterium nodatum group, Catonella, Aggregatibacter, Butyrivibrio, Lautropia, |
| **Oxidation** | **Homocysteine** | Eubacterium nodatum group, Bacteroides | Prevotella, Lautropia, Aggregatibacter, Centipeda, Filifactor, Mycoplasma, Peptococcus |

**Supplementary Table 1.** Oral bacterial genera showing positive or negative correlation with glucose, lipids, CRP, hbA1c, and oxidation.

### 2. Differential Abundance analysis of the Gut Microbiome and Saliva Microbiome in CI and NC patients

| | | | **Increased abundance** | **Decreased abundance** |
|---|---|---|---|---|
| **Cognition CI vs NC** | **Stool** | Phylum | | *Firmicutes* |
| | | Class | | *Clostridia* |
| | | Order | | *Oscillospirales, Erysipelotrichales* |
| | | Family | | *Erysipelatoclostridiaceae, [Eubacterium] coprostanoligenes group, Oscillospiraceae* |
| | | Genus | | *[Eubacterium] brachy group, Family XIII UCG-001, Holdemania, Shuttleworthia, Subdoligranulum, UCG-005* |
| | **Saliva** | Phylum | *Synergistota, Fusobacteriota* | |
| | | Class | *Synergistia, Fusobacteriia* | |
| | | Order | *Synergistales, Mycoplasmatales, Clostridia UCG-014, Fusobacteriales* | |
| | | Family | *Synergistaceae, Mycoplasmataceae* | |
| | | Genus | *Dialister, Fretibacterium, Mycoplasma* | |

**Supplementary Table 2.** Summary of differential oral and gut taxa between CI and NC patients.

### 3. Enhancing the functional potential of the gut and oral microbial communities associated with cognitive health using KEGG pathways

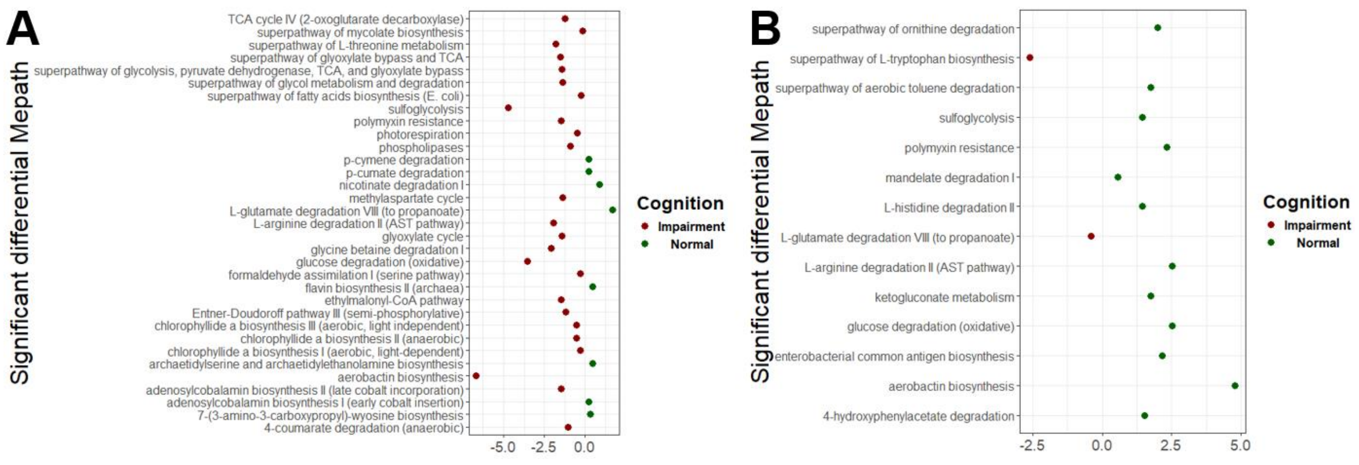


**Supplementary Figure 1.** Predicted MePath associated with cognition, comparing the Normal and Impairment groups for gut microbiome (A) and oral microbiome (B).

| | | Increased level | Decreased level |
|---|---|---|---|
| **ECs** | **Oral** | CoA-disulfide reductase; Ethanolaminephosphotransferase | Catechol 1,2-dioxygenase;D-arabinitol 4-dehydrogenase;Ferric-chelate reductase (NADPH); Tryptophan 7-halogenase; Acyl-homoserine-lactone synthase; Vanillin synthase; Trans-feruloyl-CoA hydratase; Aerobactin synthase; N(2)-citryl-N(6)-acetyl-N(6)-hydroxylysine synthase; N(6)-hydroxylysine O-acetyltransferase; Oligogalacturonide lyase; Formate dehydrogenase-N; Mannosyl-3-phosphoglycerate phosphatase; NAD(P)(+) transhydrogenase (Si-specific); Streptomycin 3-kinase; Lipid IV(A) 4-amino-4-deoxy-L-arabinosyltransferase; dTDP-4-amino-4,6-dideoxy-D-galactose acyltransferase; 1,6-dihydroxycyclohexa-2,4-diene-1-carboxylate dehydrogenase; Benzoate 1,2-dioxygenase; Sulfolactaldehyde 3-reductase; Aliphatic aldoxime dehydratase;L-lysine N(6)-monooxygenase (NADPH); 23S rRNA (guanine(1835)-N(2))-methyltransferase; Acireductone dioxygenase (Ni(2+)-requiring); Acireductone dioxygenase (Fe(2+)-requiring); Pectin lyase; TDP-N-acetylfucosamine:lipid II N-acetylfucosaminyltransferase; Formimidoylglutamate deiminase |
| | **Gut** | 4-aminobutyrate--pyruvate transaminase; Malyl-CoA lyase;(2S)-methylsuccinyl-CoA dehydrogenase;5-aminolevulinate synthase;Malonyl-CoA decarboxylase; Dimethylglycine dehydrogenase; NAD(+)--dinitrogen-reductase ADP-D-ribosyltransferase; Methylamine dehydrogenase (amicyanin); Succinyl-CoA--L-malate CoA-transferase; Precorrin-6A synthase (deacetylating); 3-methylfumaryl-CoA hydratase; Demethylspheroidene O-methyltransferase; 1-hydroxycarotenoid 3,4-desaturase; Alanine--glyoxylate transaminase; Serine--glyoxylate transaminase; Serine--pyruvate transaminase; 4-methylaminobutanoate oxidase (formaldehyde-forming); Salicylate 1-monooxygenase; 4-hydroxybenzoate--CoA ligase; Gluconate/galactonate dehydratase; Dimethyl sulfide:cytochrome c2 reductase; Dimethyl sulfide:cytochrome c2 reductase; Pyridoxamine--pyruvate transaminase; 2-methylfumaryl-CoA isomerase; | Spermidine dehydrogenase; GalNAc(5)-diNAcBac-PP-undecaprenol beta-1,3-glucosyltransferase; Acetylserotonin O-methyltransferase; Nicotinate dehydrogenase (cytochrome); Non-reducing end beta-L-arabinofuranosidase; 15,16-dihydrobiliverdin:ferredoxin oxidoreductase; dCTP diphosphatase |

**Supplementary Table 2.** Top 30 differential oral and gut ECs between CI and NC patients.

| | | Increased level | Decreased level |
|---|---|---|---|
| KOs | Oral | K16819 | K02465, K08087, K19734, K09916, K18555, K03381, K18838, K07349, K03225, K00007, K03226, K03230, K03222, K03228, K03219, K03229, K14744, K16324, K03812, K07350, K07310, K11911, K16090, K12290, K09016, K10972, K03220, K07348, K10973 |
| | Gut | K18588, K07716, K14981, K14980, K07234, K13587, K13583, K00410, K13588, K17662, K16871, K13584, K09991, K09987, K18587, K08691, K14448, K06602, K00643, K14447, K01578, K15721, K02132, K00315, K05951, K13013, K13598, K18989, K18990, K07167 | |

**Supplementary Table 3.** Top 30 differential oral and gut KOs between CI and NC patients.

#### 3.1 Stool samples

| ID | Description | BH p.adjust | KOs ID | count |
|---|---|---|---|---|
| ko00195 | Photosynthesis | 7.703e-22 | K02705, K02639, K02720, K02703, K02700, K02643, K02718, K02714, K02640, K02711, K02722, K02723, K02717, K02697, K02634, K02694, K02704, K02707, K02709, K02716, K08902, K08903, K02724, K02638, K02693, K02698, K02706, K08906, K02641 | 29 |
| ko00196 | Photosynthesis - antenna proteins | 8.927e-17 | K02290, K05378, K05379, K05381, K05383, K05384, K05386, K05385, K05377, K05382, K05376, K02092, K02094, K02096, K02097, K02288, K02289, K02093, K02095, K02284, K02285 | 21 |
| ko03070 | Bacterial secretion system | 4.318e-9 | K02465, K11915, K11016, K11017, K02461, K02462, K02460, K02458, K02459, K02457, K11912, K02452, K03223, K03229, K13301, K04059, K03221, K03425, K03226 | 19 |
| ko02026 | Biofilm formation - Escherichia coli | 6.419e-7 | K04335, K04333, K04336, K12687, K04334, K07782, K07677, K07678, K07676, K02425, K07689, K02403, K02402, K14051, K07687 | 15 |
| ko01503 | Cationic antimicrobial peptide (CAMP) resistance | 7.544e-7 | K19079, K13632, K19238, K07264, K07660, K07806, K07637, K12963, K12973, K07771, K13014, K12975, K19236, K07643 | 14 |
| ko00680 | Methane metabolism | 2.685e-6 | K08691, K00830, K01086, K13942, K00400, K13812, K00442, K00440, K00319, K00443, K00578, K00579, K00580, K00581, K00582, K00583, K00204, K07811, K10978, K01070, K03533, K03532, K18277, K00399, K13039, K06034 | 26 |
| ko01200 | Carbon metabolism | 0.000151 | K08691, K14448, K14447, K09709, K14472, K00830, K01086, K14471, K05308, K01602, K14470, K07511, K15918, K13942, K13812, K00319, K00578, K00579, K00580, K00581, K00582, K00583, K00204, K00116, K01637, K01690, K01638, K01782, K01070, K01682, K01825, K00399, K00242, K19243 | 34 |
| ko02040 | Flagellar assembly | 0.000248 | K02394, K02393, K02386, K02399, K02391, K02423, K10941, K02425, K02403, K02402, K03516 | 11 |
| ko04112 | Cell cycle - Caulobacter | 0.000514 | K07716, K13587, K13583, K13588, K13584, K13589, K06985, K13582 | 8 |
| ko00860 | Porphyrin metabolism | 0.00116 | K00643, K02228, K11336, K04037, K04039, K05369, K00214, K05370, K05371, K10960, K04040, K04038, K03428, K03403, K13543, K07215, K00228 | 17 |
| ko00984 | Steroid degradation | 0.00409 | K15981, K03333, K16049, K16048, K05898 | 5 |
| ko00630 | Glyoxylate and dicarboxylate metabolism | 0.00450 | K08691, K14448, K14447, K01432, K00830, K01602, K15918, K01637, K01638, K01682, K12972, K03418, K00090 | 13 |
| ko00190 | Oxidative phosphorylation | 0.00451 | K00410, K02132, K05575, K05577, K05586, K05588, K05578, K05572, K05573, K05581, K05583, K05584, K05585, K05582, K05579, K05574, K00242, K02297, K02300, K02298, K02299 | 21 |
| ko00364 | Fluorobenzoate degradation | 0.00829 | K01061, K05783, K05549, K05550, K05784 | 5 |
| ko00540 | Lipopolysaccharide biosynthesis | 0.0106 | K02848, K03275, K03276, K12981, K12985, K07264, K11211, K12973, K12975 | 9 |
| ko00740 | Riboflavin metabolism | 0.0158 | K08096, K14653, K19286, K03788, K01497, K05368, K12152, K00299 | 8 |
| ko00640 | Propanoate metabolism | 0.0199 | K01578, K07511, K11264, K00932, K01782, K01682, K01825, K17489, K19745, K13923, K01908 | 11 |

| | | | | |
|---|---|---|---|---|
| ko00622 | Xylene degradation | 0.0219 | K10622, K05783, K05549, K05550, K10621, K05784 | 6 |
| ko00410 | beta-Alanine metabolism | 0.0219 | K01578, K00316, K07511, K13799, K01782, K01825, K00276 | 7 |
| ko00920 | Sulfur metabolism | 0.0220 | K17225, K17285, K16964, K16965, K16966, K00988, K08353, K08354, K00381, K00299, K03119, K04091 | 12 |
| ko00130 | Ubiquinone and other terpenoid-quinone biosynthesis | 0.0220 | K18534, K12073, K18286, K03181, K18800, K19222, K03184, K03185 | 8 |
| ko00030 | Pentose phosphate pathway | 0.0266 | K01086, K05308, K13812, K01690, K00032, K11441, K05774, K00090, K00117, K19243 | 10 |
| ko02030 | Bacterial chemotaxis | 0.0284 | K03776, K05876, K12368, K05874, K10108 | 5 |
| ko00362 | Benzoate degradation | 0.0290 | K04105, K04107, K01782, K01825, K10622, K05783, K05549, K05550, K14727, K10621, K05784 | 11 |
| ko00260 | Glycine, serine and threonine metabolism | 0.0345 | K00643, K00315, K15783, K15786, K15784, K00830, K15918, K00276, K12525, K12972, K00090 | 11 |
| ko00760 | Nicotinate and nicotinamide metabolism | 0.0385 | K19191, K18028, K13995, K15357, K14974, K18030, K18029, K08324, K00322, K11751 | 10 |
| ko00240 | Pyrimidine metabolism | 0.0385 | K16904, K13799, K09023, K09021, K09018, K08723, K09024, K09019, K08320, K09913, K11751 | 11 |
| ko00660 | C5-Branched dibasic acid metabolism | 0.0385 | K08691, K18288, K14472, K14471, K18289 | 5 |

**Supplementary Table 4**. Summary of differential gut KOs grouped in their KEGG pathways.

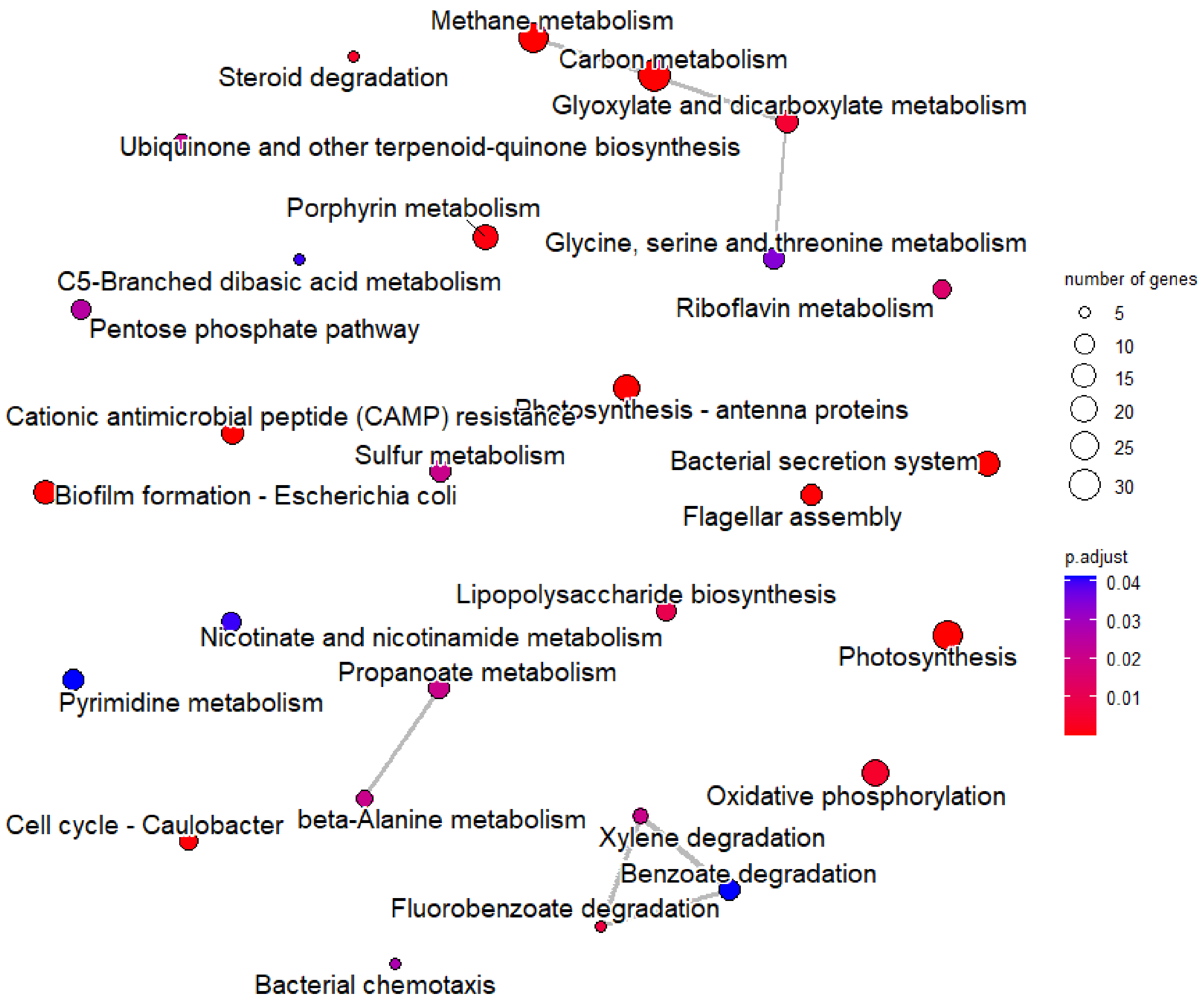

**Supplementary Figure 2.** Emap plot of enriched KEGG pathways illustrating their network association for gut microbiome.

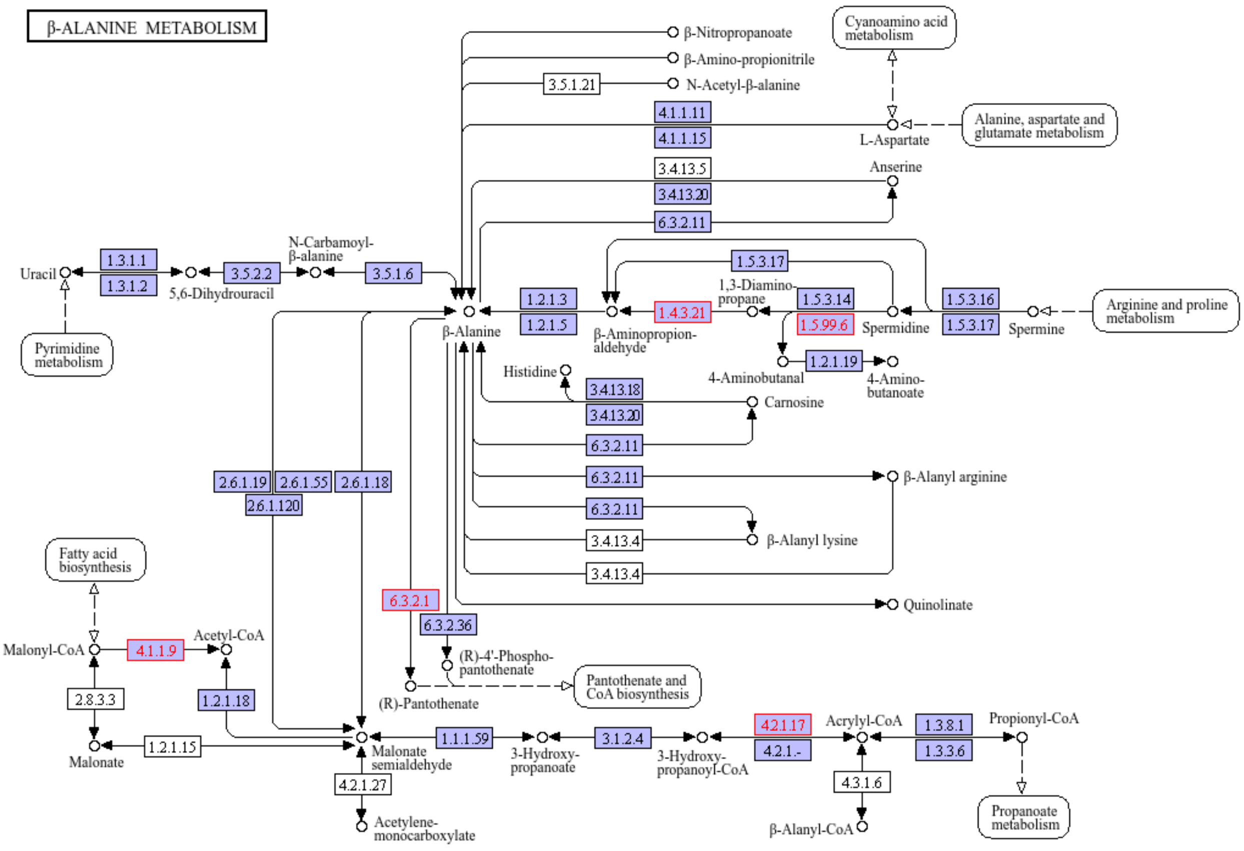


**Supplementary Figure 3.** KEGG metabolic map highlighting the KOs (red color) that were identified in *beta-Alanine metabolism*.

#### 3.2 Saliva samples

| ID | Description | BH p.adjust | KOs ID | count |
|---|---|---|---|---|
| ko03070 | Bacterial secretion system | 3.359e-12 | K02465, K03225, K03226, K03230, K03222, K03228, K03219, K03229, K03221, K02452, K02459, K11016, K02460, K02464, K02458, K02462, K02457, K02461 | 18 |
| ko01503 | Cationic antimicrobial peptide (CAMP) resistance | 7.815e-9 | K19238, K19077, K19080, K07264, K07806, K01406, K07771, K10011, K12963, K07660, K12975, K13014, K12973 | 13 |
| ko00330 | Arginine and proline metabolism | 0.00105 | K00137, K05526, K09472, K12252, K06447, K00673, K01484, K1225, K12254, K00840, K09473 | 11 |
| ko00364 | Fluorobenzoate degradation | 0.00105 | K03381, K05783, K05550, K05549, K01721 | 5 |

| | | | | |
|---|---|---|---|---|
| ko00440 | Phosphonate and phosphinate metabolism | 0.00114 | K00993, K06165, K06164, K05780, K06166, K06163, K06162 | 7 |
| ko00860 | Porphyrin metabolism | 0.001602 | K02225, K13543, K03394, K05895, K02304, K06042, K02189, K05936, K05934, K02191, K03399, K13542 | 12 |
| ko00627 | Aminobenzoate degradation | 0.00274 | K11311, K05599, K10219, K18541, K05600, K16320, K16319, K01865, K01721 | 9 |
| ko00907 | Pinene, camphor and geraniol degradation | 0.00284 | K13776, K01782, K13774, K13779, K01825 | 5 |
| ko00051 | Fructose and mannose metabolism | 0.00284 | K00007, K07026, K18334, K18335, K19270, K19291, K19290, K01795, K07046, K00045 | 10 |
| ko02026 | Biofilm formation - Escherichia coli | 0.00469 | K03567, K04333, K07689, K07677, K18968, K14051, K07687 | 7 |
| ko00350 | Tyrosine metabolism | 0.00547 | K13951, K13574, K01826, K05921, K00146, K10219, K00483, K00455 | 8 |
| ko00362 | Benzoate degradation | 0.00547 | K03381, K05783, K05550, K05549, K07823, K10219, K01782, K01825, K01055 | 9 |
| ko02025 | Biofilm formation - Pseudomonas aeruginosa | 0.00748 | K13060, K13489, K13491, K11444, K07689, K18100, K19291, K10941 | 8 |
| ko00071 | Fatty acid degradation | 0.0145 | K13951, K06445, K00529, K00255, K01782, K01825 | 6 |
| ko01240 | Biosynthesis of cofactors | 0.0145 | K02225, K03184, K13543, K18800, K03394, K05895, K02304, K19222, K01432, K06042, K02189, K05936, K05934, K02191, K03399, K13940, K13542, K06989 | 18 |
| ko01220 | Degradation of aromatic compounds | 0.0227 | K03381, K05783, K05550, K05549, K01826, K00529, K05921, K10219, K00483, K00455, K10676, K01055 | 12 |
| ko00543 | Exopolysaccharide biosynthesis | 0.0371 | K12582, K16701, K02852, K03819, K19291, K19290 | 6 |

**Supplementary Table 5**. Summary of differential oral KOs grouped in their KEGG pathways.

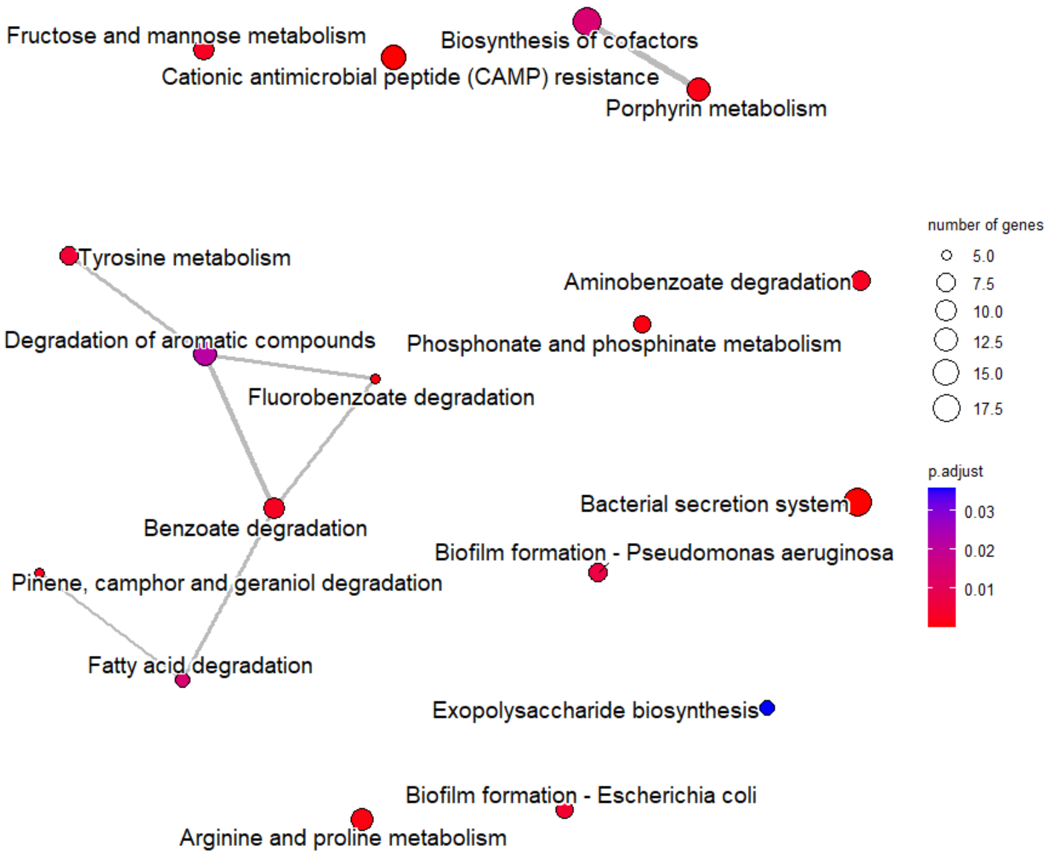


**Supplementary Figure 4.** Emap plot of enriched KEGG pathways illustrating their network association for oral microbiome.

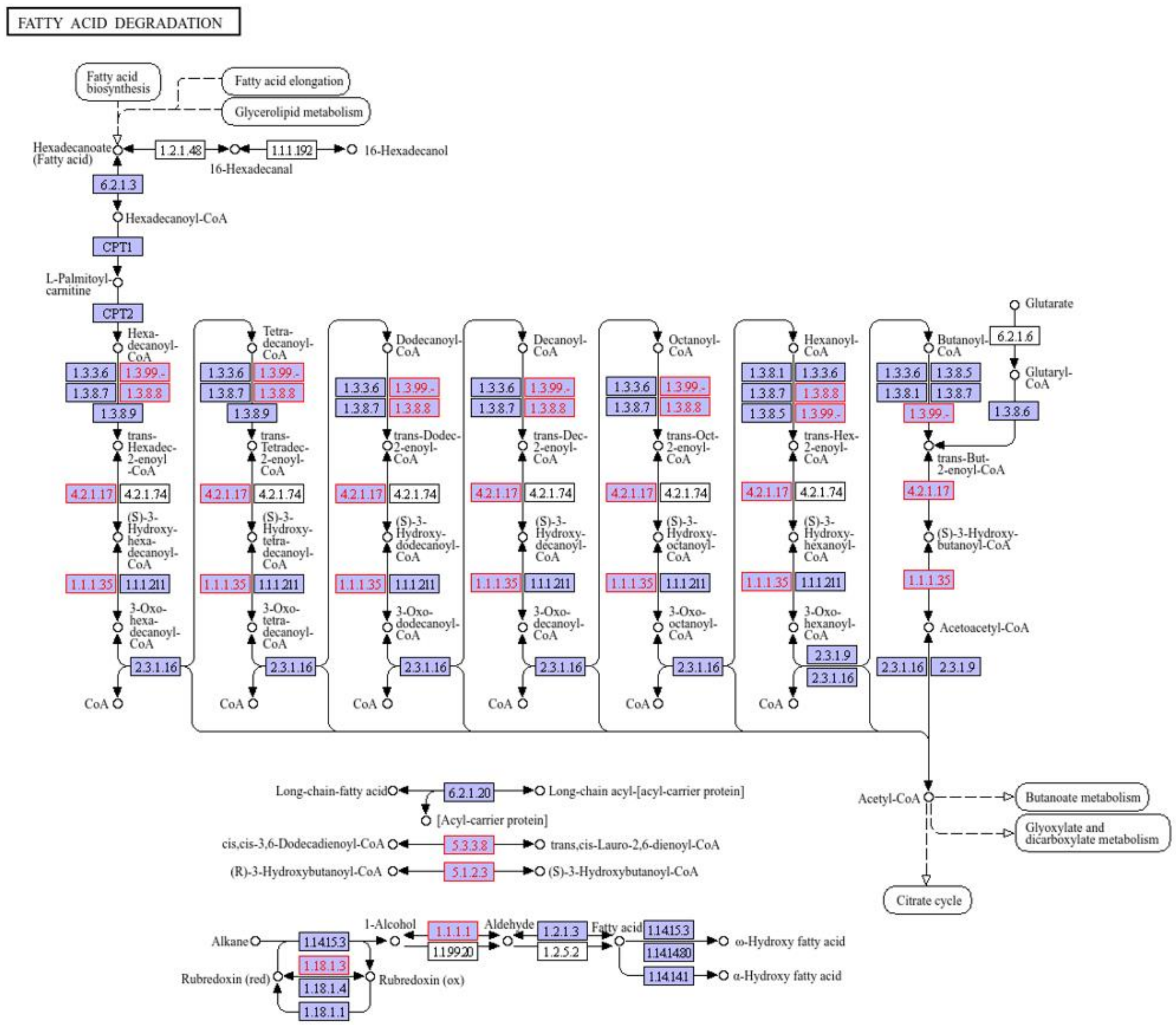


**Supplementary Figure 5.** KEGG metabolic map highlighting the KOs (red color) that were identified in *Fatty acid degradation*.
